# Xenon Explores Apparent and Cryptic Binding Sites

**DOI:** 10.1101/2025.04.30.651394

**Authors:** Shinji Iida

## Abstract

Molecular binding sites of a protein tell us how to target the protein from their structural information, which is required for structure-based drug design. However, regardless of their importance, some binding sites, called cryptic binding (CB) sites, are not always readily apparent in the absence of ligands. The identification of CB sites from protein structures remains challenging, and no standard method has been proposed for this purpose. In this study, inspired by the observation of X-ray protein structures with noble gases, particularly xenon, I evaluate a xenon-based CB site identification strategy from their apo states by directly comparing its performance against benzene as a baseline probe. I performed atomistic molecular dynamics simulations of apo proteins with known CB sites in the presence of xenons. The simulations have revealed that xenons are likely to occupy not only apparent binding sites but also CB sites, implying that they serve as a probe to explore binding sites.

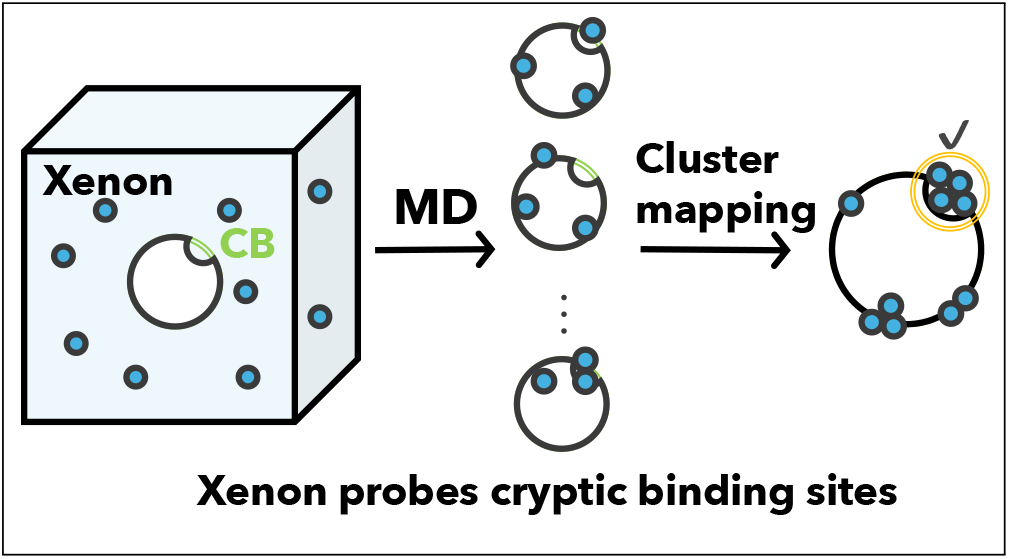

## Introduction

Proteins adopt specific three-dimensional structures that enable them to interact with other molecules, playing essential roles in biological processes. These interactions often involve concave regions on the protein surface, commonly referred to as pockets or binding sites. Accordingly, various binding site detection algorithms have been developed to provide structural insights relevant to drug design. ^1–3^

However, applying such algorithms to a single, static structure risks overlooking transiently formed binding sites. In reality, some binding sites are not visible in the unbound (apo) state but become apparent upon ligand binding (holo state). Such sites are known as cryptic binding (CB) sites?a term first introduced, to my knowledge, by Frembgen-Kesner and Elcock.^4^ CB sites were later comprehensively studied by Cimermancic et al., who created the dataset, CryptoSite, and trained a classical machine learning model (e.g., support vector machine) predicting that CB sites are widespread across the human proteome. ^5^

To account for the protein flexibility that reveals CB sites, several computational methods have emerged. While recent machine learning models show promise in predicting CB sites, ^6,7^ an alternative, physics-based approach involves explicitly sampling a protein’s conformational ensemble. Atomistic molecular dynamics (MD) simulations are a powerful tool for generating such ensembles, sampling structures according to the Boltzmann distribution. ^8,9^ Iida et al. proposed the CB site index, which quantifies the free energy difference associated with the solvent-accessible surface area (SASA) of an amino acid residue in its buried versus exposed state. ^9^ More recently, Meller et al. demonstrated that MD simulations initiated from AlphaFold2-generated structures can effectively sample CB site openings. ^10^

In addition to such geometric or energetic criteria, probe molecules have been employed in MD simulations to identify CB sites. ^11,12^ Yanagisawa et al. developed a rational and systematic cosolvent set for this purpose. ^11^, and more recently, Koseki et al. and Motono et al. from the same team proposed the persistent homology method ^13^ and mixed-solvent MD combined with machine learning. ^14^ Smith and Carlson applied MixMD protocols and successfully mapped known CB sites. ^12^ One of the common probes is benzene which is shown to identify ligand binding sites. ^15–17^

Among other small non-polar probes, a noble gas xenon shows the potential to detect CB sites due to its ability to bind to hydrophobic sites on a protein. ^18^ Xenon has indeed be identified as a promising probe by Vithani et al. who used it in combination with enhanced sampling simulations to explore CB sites in KRAS. ^19^ Xenon-protein interactions have been studied via X-ray crystallography and NMR spectroscopy ^20,21^ and its physiological properties. ^22,23^ Even though there were pioneering works for the investigation of the interaction of xenon with myoglobin ^24^ and an engineered mutant of T4 lysozyme, ^25^ xenon-mediated interaction for ligand binding sites has not been well studied yet.

In this study, I characterise xenon-protein interactions with atomistic MD simulations by exploring apparent and CB sites on proteins. I first built a xenon model with appropriate Lennard-Jones parameters for the reproduction of xenon’s physical properties. To assess its validity, I performed MD simulations for proteins that are known to be in complex with xenon. I then evaluate the ability of xenon to bind to apparent binding sites and CB sites for target proteins, compared with benzene molecules. Lastly, I discuss the limitations of xenon for the identification of CB sites.

## Materials and Methods

### Xenon’s model

Lennard-Jones (LJ) potential is expressed as

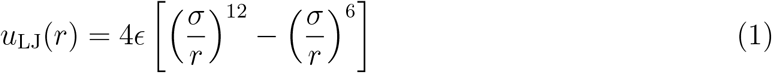

where *r* is the distance between two particles, *σ* is the distance at which the potential becomes zero (a.k.a the size of the particle), and *ϵ* is the depth of the potential well. Truncated and shifted Lennard-Jones (TSLJ) potential is then defined as

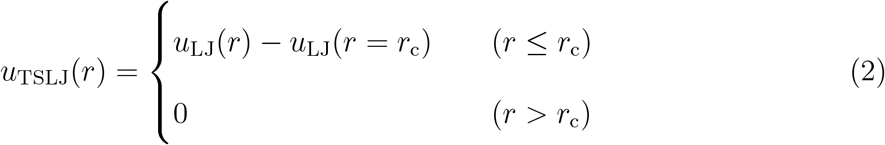

where at the cut-off distance *r*_c_, the potential becomes zero. The use of TSLJ potential avoids the calculation of long-range correction, which may be computationally demanding. ^26^ TSLJ potential is implemented in Gromacs wherein the Verlet cut-off scheme, the potential is shifted by a constant such that it is zero at the cut-off distance. ^27^ In this study, the parameters *σ* and *ϵ* of TSLJ potential were adopted from the literature where it was optimised to reproduce the vapour pressure and saturated liquid density (Table 1).^26^ The parameters are slightly different from those in the work by Tang and Toennies.^28^

**Table 1:** LJ parameters of xenon.

| $\sigma/\text{nm}$ | $\epsilon/\text{kJ mol}^{-1}$ |
| --- | --- |
| 0.39460 | 2.28531 |

One might think that xenon is a notably larger noble gas atom compared to neon, argon, and krypton and has polarisability that enables xenon to be susceptible to charge-induced dipole interaction, dipole-induced dipole interaction, and London dispersion interaction with a protein; this description aligns with reality. Nonetheless, if the xenon model expressed by the LJ parameters has successfully reproduced xenon-protein complex structures, it is reasonable to state that the model describes a crucial aspect of xenon-protein interactions.

Even though the LJ parameters correctly reproduce some physical quantities for purely xenon solution, ^26^ this does not guarantee that the LJ parameters correctly model the interactions of xenon with proteins. It is therefore necessary to confirm whether xenon with the LJ parameters binds to xenon-binding sites experimentally verified, and, in the result section, the xenon model was validated by MD simulations for five proteins known to bind to xenon. Their PDB IDs of the five proteins are as follows: 1c10^18^(lysozyme), 1c1m ^18^(elastase), and 1c3l^18^ (subtilisin carlsberg), 5hw1^29^(beta-lactamase), and 4nxa ^30^(myoglobin).

### Pairs of apo and holo PDBs

Table 2 shows the pairs of apo and holo PDB IDs. The pairs were used to define each set of amino-acid residues for binding sites, apparent binding sites, and CB sites. The pairs were adopted from the previous work, ^9^ being selected such that the ligand binding affinity (IC50 or *K*_d_) is in the order of nM in the CryptoSite dataset. ^5^

**Table 2:** Pairs of apo and holo structures.

| Pair |  | Pair |  | Pair |  | Pair |  |
| --- | --- | --- | --- | --- | --- | --- | --- |
| Apo | Holo | Apo | Holo | Apo | Holo | Apo | Holo |
| 1b6b<sup>31</sup> | 1kuv<sup>32</sup> | 1bsq<sup>33</sup> | 1gx8<sup>34</sup> | 1fvr<sup>35</sup> | 2oo8<sup>36</sup> | 3cj0<sup>37</sup> | 2brl<sup>38</sup> |
| 1hag<sup>39</sup> | 1ghy<sup>39</sup> | 1jbu<sup>40</sup> | 1wun<sup>41</sup> | 1k5h<sup>42</sup> | 2egh<sup>43</sup> | 2ohg<sup>44</sup> | 2ohv<sup>44</sup> |
| 1m47<sup>45</sup> | 1py2<sup>46</sup> | 1maz<sup>47</sup> | 2yxj<sup>48</sup> | 1nep<sup>49</sup> | 2hka<sup>49</sup> | 3f74<sup>50</sup> | 3bqm<sup>51</sup> |
| 1uk3<sup>52</sup> | 2gz7<sup>53</sup> | 1w50<sup>54</sup> | 3ixj<sup>55</sup> | 2wgb<sup>56</sup> | 2v57<sup>56</sup> | 5j2v<sup>57</sup> | 2wi7<sup>58</sup> |

### Definition of relative solvent accessible surface area

Solvent accessible surface area (SASA) is defined as the surface area traced with a probe whose radius is set to 1.4 Å. Relative SASA was defined for each amino-acid residue by

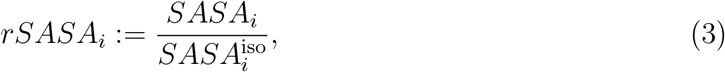

where *SASA*_*i*_ is for an *i*-th amino-acid residue in a protein, and 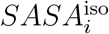is for an iso-lated amino-acid residue with its two nearest neighbours if present. It should be noted that *rSASA*_*i*_ ranges from 0 to 1 where “0” corresponds to a completely buried state and “1” does to a completely exposed state. The *rSASA*_*i*_ was computed by the function *cmd*.*get sasa relative* implemented in PyMOL ^59^

### Definition of binding sites

Let us define a set of contact amino-acid residues as a binding site. Suppose that apo structure and its corresponding holo structures are given. Let *P* and *L* be a set of aminoacid residues in a protein and a set of atoms in the ligand bound to the target protein, respectively. With the cutoff distance *d*_c_ = 5 Å, the binding site of the molecule was defined by a set of contact residues:

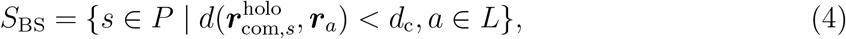

where 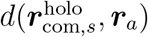 is the distance between the centre-of-mass coordinates 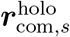of the sidechain for an amino-acid residue *s* in the holo structure and the coordinates ***r***_*a*_ of an atom *a*in *L*. The cutoff distance was set to the sum of typical Van der Waals radii (2.0 Å), yielding5.0 Å, with an additional 1 Å buffer.

### Definition of apparent and cryptic binding sites

Here, apparent and CB sites were defined according to an apo state and its corresponding holo state (Figure 1). The definition was used for evaluation of probe-binding events.

**Figure 1:**
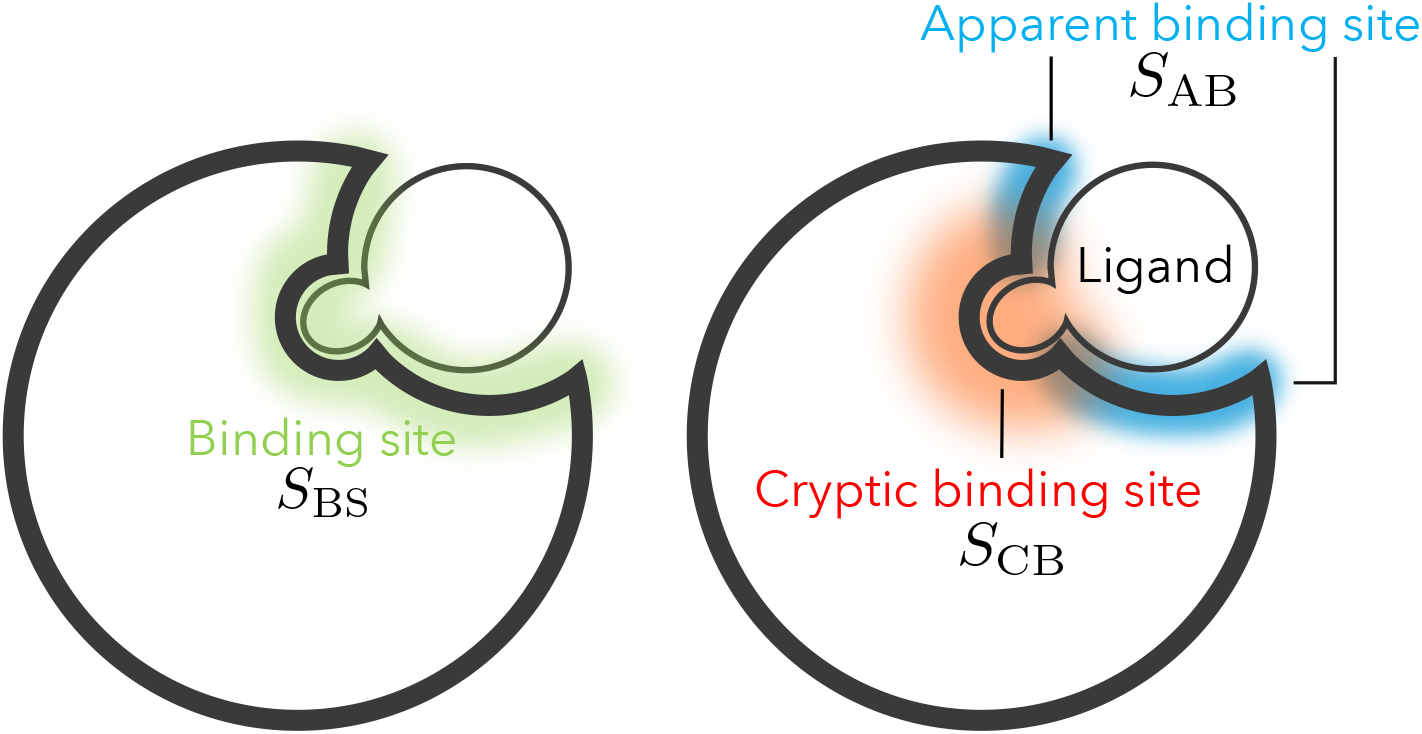
A binding site consists of two regions: apparent and CB sites. The left panel shows a binding site bound to a ligand, while the right panel shows the two regions, an apparent and a CB site, in the binding site. Apparent and CB sites are coloured by blue and orange, respectively. Since the definition of CB sites in this study was based on changes in solvent exposure (Δ*rSASA*), it enables a clear distinction between CB sites and apparent binding sites.

The *rSASA*_*i*_ of each residue in *S*_BS_ was calculated for the apo structure and the ligand-free holo structure (i.e., the ligand was removed but the protein structure was kept), and then the difference is given by

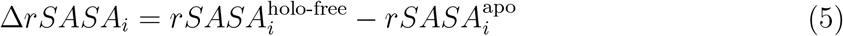

For a tolerance of exposure *ϵ*, if Δ*rSASA*_*i*_ > *ϵ*, then the *i*-th residue was considered to be in a CB site. Namely, a CB site was defined by

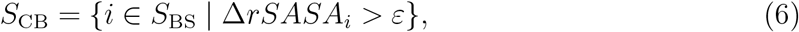

Because Δ*rSASA*_*i*_ > 0 indicates that the *i*-th residue is exposed in the holo structure but is less exposed in the apo structure, it represents an essential aspect of CB sites. The tolerance *ϵ* was set to 0.1 in this study. The set of residues in a CB site were provided in Table S1.

Now, the apparent binding site was defined by a set of contact amino-acid residues except for those in a CB site (Table S2):

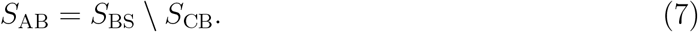

### Molecular dynamics simulation

For the validation of xenon model, five independent 100-ns NPT simulations at 300 K of each known xenon-binding protein (PDB IDs: 1c10, ^18^ 1c1m,^18^ 1c3l, ^18^ 5hw1,^29^ and 4nxa ^30^) were performed in the presence of xenon. Note that the simulations were initiated from a protein structure where xenons are not bound. For two systems (PDB IDs: 1c10 and 5hw1), it was observed that xenon atoms rarely bound to the known xenon-binding sites within 100 ns × 5 runs even though xenon should diffuse internal cavities. To enhance the diffusion, five independent 100-ns NPT simulations at 400 K were also carried out for the two systems, being checked if xenons correctly find the known xenon-binding sites. For each of the sixteen protein systems with CB sites (Table 2), five independent 100-ns MD simulations were performed in the presence of xenon.

The following procedure was used to set up each system: (1) protonation states of histidine residues were assigned using *pdb2gmx* command. (2) The resultant PDB files were processed with *tleap*, generating topology and coordinate files in Amber format (i.e., .prmptop and .inpcrd). Disulphide bonds were correctly formed in this step. (3) These Amber-format files were converted to Gromacs format (i.e., .top and .gro) using ParmEd ^60^ (https://github.com/ParmEd/ParmEd). (4) These systems were resolvated with dodecahedron water box using *gmx* tools. (5) Xenons (or benzenes) were inserted using *gmx insert-molecule*, replacing some water molecules; the concentration of xenons (or benzenes) was set to 0.1M. Although this concentration was not systematically optimised, Figures S1 and S2 indicated that the concentration did not destabilise protein structures, showing that 0.1 M was deemed a reasonable compromise. (6) Ions were added to neutralise the system. (7) The system, including protein, water, ions, and xenons (or benzenes), was energy-minimised with positional restraints on protein backbones, then equilibrated using NVT and NPT ensembles (500 ps each). (8) Five independent production runs of 100 ns each were then performed for each system under NPT ensemble.

Computational settings were as follows: Time step was set to 2 fs, and an interaction table for cut-off scheme ^61^ was used whose list was updated every 20 fs. The leap-frog algorithm was used for an integrator, and LINCS ^62^ was for hydrogen-bond constraints (lincs-iter=1, lincs-order=4). V-rescale ^63^ was for temperature control for two groups, protein and nonprotein (*τ*_*T*_ = 0.1, compressibility = 4.5 × 10^6^). Berendsen pressure coupling ^64^ was used for the first box equilibration, and Parrinelo-Rahman coupling for all production runs to keep 1 atm pressure (*τ*_*p*_ = 2.0, compressibility = 4.5 × 10^6^). Short-range electrostatic and Van der Waals interaction cut-off values were set to 10 Å. Amber ff14SB ^65^ was employed for protein force field and TIP3P model ^66^ for water molecules. The particle-mesh Ewald (PME) method^67^ was used for long range interactions calculation. Snapshots were stored for every 10 ps. Gromacs 2022.4^68^ was used for all simulations. The force field parameters of benzene were assigned with GAFF2 via Acpype. ^69^

The initial atomic coordinates for myoglobin, which forms a xenon-protein complex, were obtained from the X-ray crystal structure (PDB ID: 4nxa). ^30^ The protein chain was isolated, and the protonation states of ionisable residues were determined at a physiological pH using the H++ web server. ^70^ Force field parameters for the non-standard active site comprising a heme were generated using the Metal Center Parameter Builder (*MCPB*.*py*) program^71^ within the AmberTools23 suite. ^72^ This process involved a multi-step quantum mechanical (QM) approach. Two distinct models of the active site were constructed for the QM calculations: A small model, used to derive bonded parameters (bond and angle force constants), consisted of the heme group, the Fe^3+^ ion, the coordinating water molecule, and the imidazole ring of the proximal histidine. A large model, used to derive partial atomic charges, included the same components as the small model, but the proximal histidine was capped with acetyl and N-methylamide groups to neutralise the termini and accurately represent the peptide backbone’s electronic environment.

Initial parameter files (frcmod) for the constituent heme, Fe^3+^, and water molecules were generated with the *parmchk2* tool. All QM calculations were performed with the Gaussian 16 software package ^73^ using the B3LYP density functional ^74^ and the 6-31G* basis set. The geometry of the small model was fully optimised, and a subsequent frequency calculation was performed on the optimised structure. The resulting Hessian matrix was used to derive the force constants for the bonds and angles associated with the metal centre. A separate single-point QM calculation was performed on the large model to generate the electrostatic potential (ESP). Restrained Electrostatic Potential (RESP) fitting was then used to assign partial atomic charges to all atoms in the large model. Finally, the derived bonded parameters and RESP charges were integrated to build a complete force field for the myoglobin complex. The resulting AMBER-formatted parameter and topology files were converted for use in GROMACS with ParmEd. ^60^ The MD simulation protocol was the same as that of the other systems.

### Definition of xenon clusters on protein via HDBSCAN

The clusters of xenons (or benzenes) were identified by HDBSCAN (Hierarchical Density-Based Spatial Clustering of Applications with Noise), which can detects clusters with different densities and an arbitrary shape and removes outliers. ^75^ Below is the procedure of clustering:

1. From MD trajectories, the snapshots of xenon or benzene were extracted and then flattened, so that a single, aggregated coordinates were obtained.
2. HDBSCAN was applied to the single structure, generating xenon/benzene clusters *c*_*i*_ (*i* = 1, 2, …, *N*_clusters_). The only parameter optimised was *min clust size* which was set to the number of xenons in the single structure weighted by 0.0025. This weight was selected by the elbow method (See Figures S3 and S4 in the supporting information). The other parameters were set to the default values (See Table S3).

### Hit rate and false discovery rate of apparent and cryptic binding sites

To provide an assessment of the ability of xenon/benzene to apparent and CB sites, two complementary metrics were used: the hit rate (also known as recall or true positive rate) and false discovery rate (FDR). This pair of metrics is well-suited for binding site identification, which represents a classification problem on a highly imbalanced dataset where the number of positive sites is vastly smaller than the negative sites of the protein.

Hit rate was defined by the following steps:

1. The coordinates of probes (i.e., xenon/benzene) obtained from MD simulations were clustered as described in the previous section. The clusters *c*_*i*_ (*i* = 1, 2, …, *N*_clusters_) were sorted in an descending order based on the number of points in each cluster. The largest cluster was considered to be the most plausible one. The top *n*_top_ clusters were used for further evaluation.
2. The top *n*_top_ clusters were mapped onto the corresponding apo structure, which was aligned with the MD snapshots beforehand.
3. For each amino-acid residue in the apo structure, if its centre-of-mass point 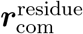 was within 4 Å of the closest atom of a probe in any of the top *n*_top_ clusters, then the residue was considered to be probe-binding, and the set of the probe-interacting residues is denoted as *S*_*C*_. The use of only the top one cluster (*n*_top_ = 1) to identify probe-interacting residues and determine binding site identification is the most stringent approach, as it limits the spatial regions considered to only the largest and most probable aggregation of probe atoms identified by the clustering method.
4. If there is any overlap between *S*_*C*_ and a predefined set of apparent *S*_AB_ or CB site residues *S*_CB_ (i.e., *S*_*C*_ ∩ *S*_type_ ≠ ∅, type = apparent or cryptic), then the xenons were considered to be bindable (*hit*/true positive) to an apparent or CB sites.
5. Step 1 - 4 was repeated for each system. The overall hit rate was then calculated by dividing the total number of hits by the total number of systems.

False discovery rate (FDR) is a useful metric to evaluate classification performance on imbalanced datasets. This is relevant to the present study because the number of true binding sites is much smaller than the number of other potential to which xenon or benzene can bind. FDR is defined as the proportion of false positives among all predictions, and is also related to precision (= 1 − FDR). In this study, the total number of predictions correspond to *n*_top_ × *N*_sys_ (*N*_sys_ is the number of systems), as the top *n*_top_ clusters for each of the *N*_sys_ systems was analysed. A cluster is classified as a false positive (FP) if it identifies a pocket that is not a known apparent binding or CB site. FDR is then expressed by 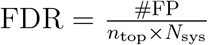 where *N*_sys_ = 16).

### Free energy mapping of xenon positions onto protein

The evaluation of the xenon model requires the identification of free-energetically stable sites on a protein. For this, the density *ρ*(***x***) of xenon around a protein within a voxel was computed with the *Density object* of [MDAnalysis](https://docs.mdanalysis.org/stable/documentation_pages/analysis/density.html#density-object).^76,77^ The free energy was computed by

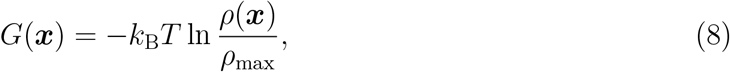

where ***x*** ∈ ℝ^3^ denotes Cartesian coordinates in a simulation box and *k*_B_ and *T* are Boltzmann constant and temperature, respectively. The division by the maximum density *ρ*_max_ sets the most stable voxel to be zero, and thus *G*(***x***) ≥ 0. It should be noted that each MD frame was superimposed before this analysis.

## Results and discussion

### Xenon model reproduces xenon-protein complex structures

The xenon model must reproduce xenon-protein complex structures. In order to verify the quality of the xenon model, I performed MD simulations of five proteins that are known to have xenon-binding sites. Mapping the free energy iso-surfaces onto protein structures, I visualised probable xenon-binding sites (Figure 2) and counted the number of xenon-binding sites that were free-energetically stable sites (Table 3). It was found that, at 300 K, xenon-binding sites overlapped with free energetically stable sites (See check marks in Figure 2). However, some xenon-binding sites for the proteins of PDB ID 1c10 and 5hw1 did not accommodate xenon atoms (See cross marks in Figure 2), because the sites are buried inside the proteins and were unreachable during the simulations.

**Table 3:** Counts of known xenon-binding sites (*n*_xs_) and the known sites corresponding to free-energetically stable xenon-binding sites (*n*_s_).

| PDB ID | $n_{\text{s}}/n_{\text{xs}}$ |
| --- | --- |
| 1c10<sup>18</sup> | 1/2 |
| 1c10 (400K) | 2/2 |
| 1c1m<sup>18</sup> | 1/1 |
| 1c3l<sup>18</sup> | 1/1 |
| 5hw1<sup>29</sup> | 1/3 |
| 5hw1 (400K) | 3/3 |
| 4nxa<sup>30</sup> | 4/5 |

**Figure 2:**
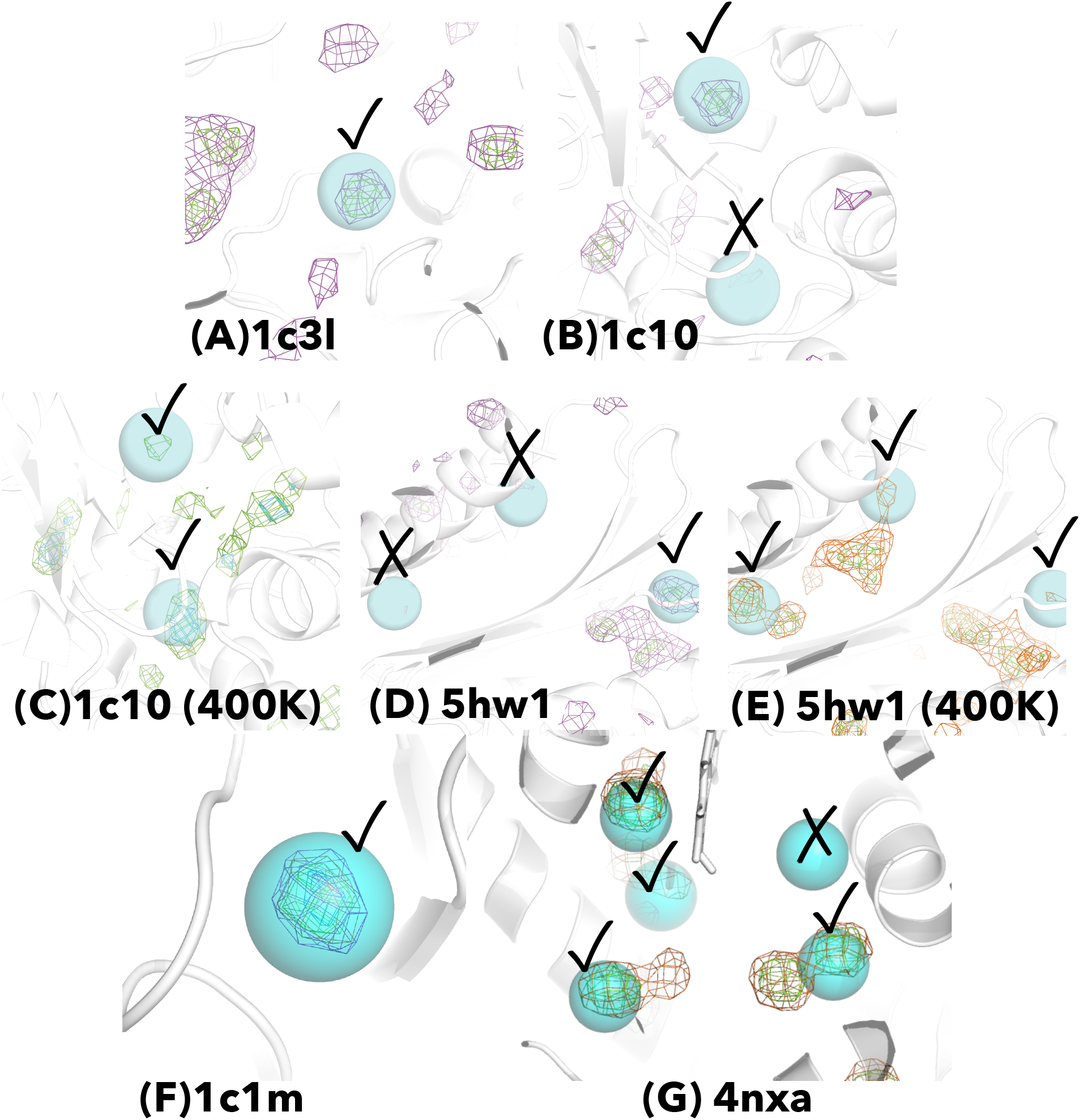
Free energy iso-surface of xenons on each target protein. This analysis was performed for a trajectory aggregated from the five trajectories of each system. Each cyan sphere represents a xenon atom in a crystal structure. Each mesh surface indicates iso-free energy surface of *F* (***x***)*/k*_B_*T* : 4.0 for magenta, 3.0 for orange, 2.0 for green, and 1.0 for cyan. The cross and check marks indicate non-explored xenon-binding sites and explored sites with high probability, respectively.

To make such buried sites reachable during short MD simulations, I also performed MD simulations at 400 K. It was found that xenon atoms were also stabilised in the sites unexplored at 300K. Moreover, even though xenon interacted with proteins nonspecifically, some of the most free-energetically stable sites corresponded to the known xenon-binding site in each X-ray structure (Figure 2). I also confirmed the reproducibility of the free energy iso-surface for the five independent MD simulation for each system (Figure S5). These results demonstrate the reproducibility of the model for xenon-protein interactions.

One may think that the simulations at 400 K caused the disruption of the whole structure, leading to the artefact that xenon appeared to be located in the xenon-binding sites; however, since the whole structure of each protein was retained during the simulations, such artefact did not occur (Figure S6).

### Xenon binds to apparent binding sites

With the xenon model, I evaluated the ability of xenon to bind to apparent binding sites with the baseline of benzene. Table 4 demonstrates that xenon and benzene can bind to apparent binding sites. If the hit rate was evaluated only for the top 1 cluster, xenon probed apparent binding sites for four out of sixteen target proteins and benzene did for two out of sixteen.

**Table 4:** Hit rate and FDR of apparent binding sites. The hit systems are shown in Tables S4 and S5.

| Used top cluster ID | Hit rate |  | FDR |  |
| --- | --- | --- | --- | --- |
|  | Xenon | Benzene | Xenon | Benzene |
| 1 | 4/16 | 2/16 | 0.75 | 0.88 |
| $\leq 2$ | 6/16 | 3/16 | 0.81 | 0.91 |
| $\leq 3$ | 9/16 | 4/16 | 0.79 | 0.92 |
| $\leq 4$ | 10/16 | 6/16 | 0.83 | 0.91 |
| $\leq 5$ | 12/16 | 6/16 | 0.86 | 0.93 |
| $\leq 6$ | 12/16 | 6/16 | 0.89 | 0.94 |
| $\leq 7$ | 12/16 | 7/16 | 0.90 | 0.94 |
| $\leq 8$ | 12/16 | 8/16 | 0.91 | 0.94 |
| $\leq 9$ | 14/16 | 8/16 | 0.91 | 0.94 |
| $\leq 10$ | 14/16 | 9/16 | 0.92 | 0.94 |

As more clusters were considered, the hit rate increased for both xenon and benzene. This suggests that (i) xenon and benzene can bind to apparent binding sites. and that (ii) xenon are more likely to bind to apparent binding sites than benzene. In fact, earlier studies also suggested that xenon is capable of binding to hydrophobic pockets. ^18,20^ The table clearly demonstrates that an apparent binding site was probed by xenon, and this result coincides with those earlier works. To my knowledge, this work offers one of the first comparative evaluations of xenon’s binding to apparent sites versus the baseline benzene.

### Xenon binds to cryptic binding sites

Similar to the analysis in the previous section, I examined the ability of xenon to bind to CB sites as a comparison with the baseline benzene. Table 5 shows that if the top 1 cluster was only considered, xenon probed CB sites of four target proteins out of sixteen, whereas benzene probed five target proteins out of sixteen. With the top 1 cluster, benzene was thus superior to xenon; however, if top ≤ 2 clusters were used for the assessment of binding, the hit rate of xenon exceeded that of benzene. As compared with Table 4, benzene struggled to bind to CB sites. For example, Figure S7 demonstrates a success and a failed cases of CB site probing by benzene. Any benzene molecules approached the ligand binding site because of the rid of the loop. While benzene molecules struggled to probe the CB site, xenon probed the CB site because of the ability to easily diffuse internal cavities (Figure S7B,C). It implies that xenon would be a better probe to identify CB sites than benzene.

**Table 5:** Hit rate and FDR of CB sites. The hit systems are shown in Tables S6 and S7.

| Used top cluster ID | Hit rate |  | FDR |  |
| --- | --- | --- | --- | --- |
|  | Xenon | Benzene | Xenon | Benzene |
| 1 | 4/16 | 5/16 | 0.75 | 0.69 |
| $\leq 2$ | 7/16 | 5/16 | 0.78 | 0.84 |
| $\leq 3$ | 10/16 | 5/16 | 0.79 | 0.90 |
| $\leq 4$ | 11/16 | 5/16 | 0.83 | 0.92 |
| $\leq 5$ | 11/16 | 5/16 | 0.86 | 0.94 |
| $\leq 6$ | 11/16 | 5/16 | 0.89 | 0.95 |
| $\leq 7$ | 11/16 | 5/16 | 0.90 | 0.96 |
| $\leq 8$ | 11/16 | 5/16 | 0.91 | 0.96 |
| $\leq 9$ | 12/16 | 6/16 | 0.92 | 0.96 |
| $\leq 10$ | 12/16 | 6/16 | 0.93 | 0.96 |

I visualised the clusters mapped onto holo structures in Figure 3. This figure shows that xenon clusters overlapped with ligand binding sites, visually indicating that xenon is able to explore CB sites.

**Figure 3:**
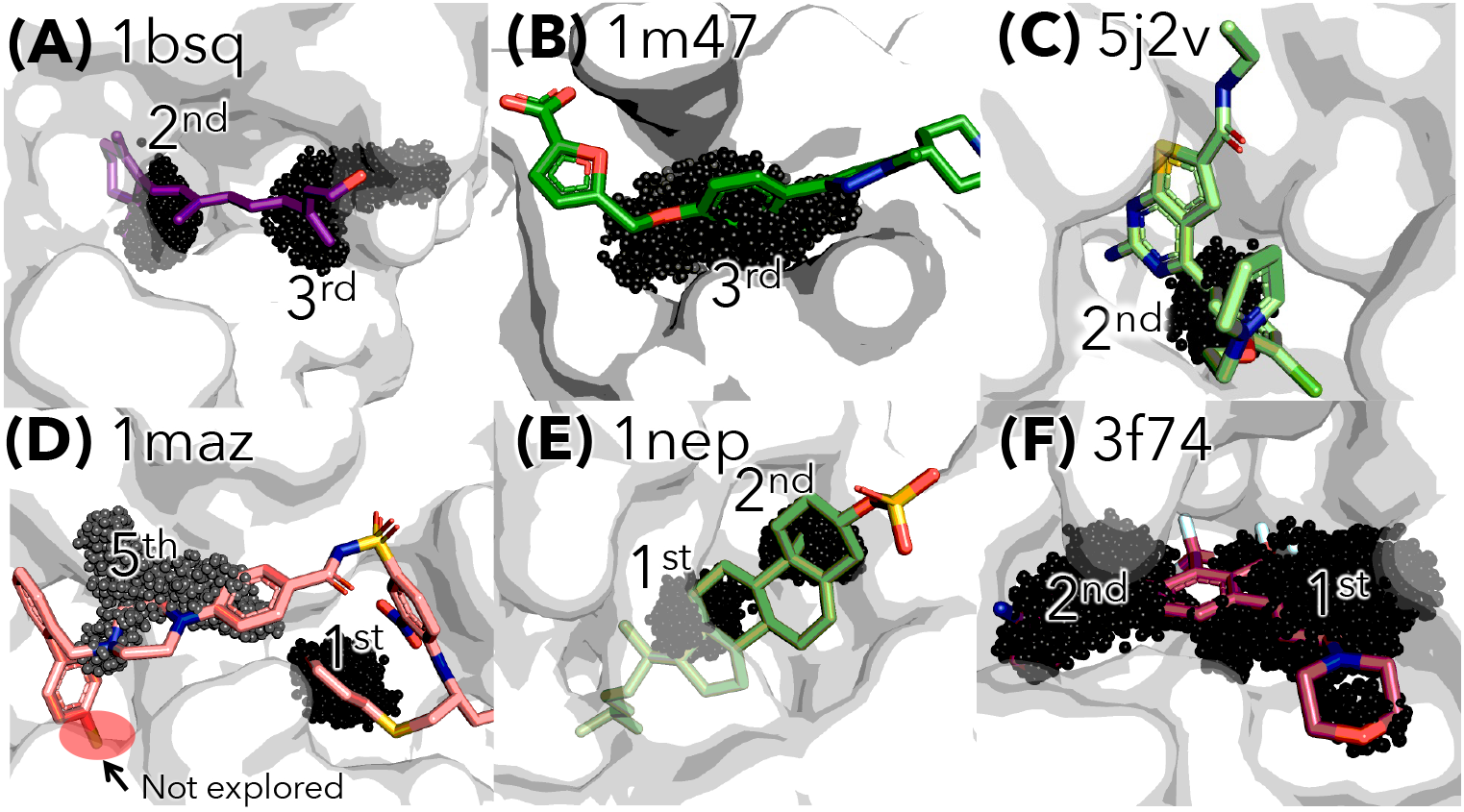
HDBSCAN clusters of xenon atoms mapped onto each holo structure corresponding to its apo structure whose PDB ID is written next to the Figure label (A)-(F). The clusters of small black spheres represent DBSCAN clusters of xenon on which cluster ID is labelled. The cases are success examples.

Let us now examine each figure in detail. In Figure 3A, the 2nd, and 3rd xenon clusters overlapped with the entire ligand. In Figure 3B, it is found that the exposure of amino-acid residues opens the CB site, and the site became accessible to xenon atoms in the 3rd cluster. In Figure 3C, the aromatic ring was traced by the 2nd xenon cluster. In Figure 3D, while the site where fluorine of the tip of the ligand is located was not probed by xenon (labelled by “Not explored”), the 5th and 1st cluster covered some aromatic rings. In Figure 3E, the four rings were probed by the 1st and 2nd clusters. In Figure 3F, the overall ligand conformation was probed by the 1st and 2nd clusters. Noting that all the simulations in this study were started from an apo structure, this result would ensure that xenon can probe CB sites even when only an apo structure is known.

While the cases above are success examples, failure cases would also be informative to point out limitations, which is discussed in the section Limitation.

### Preference of xenon binding site

Since xenon is hydrophobic, it is expected to interact with hydrophobic residues, but how about the hydrophilic residues? To investigate the ability of xenon to interact with hydrophilic residues, I analysed (i) the electrostatic surface of the binding site in each protein and (ii) contact frequency of xenon atoms to each amino-acid residue in each protein system.

Using Adaptive Poisson-Boltzmann Solver (APBS 3.4.1) in PyMOL, ^78^ I visualised the electrostatic surface (Figure 4). I found that xenon clusters are able to associate with highlycharged surfaces except for Figure 4D-F, whose surfaces are more hydrophobic than those of 4A-C.

**Figure 4:**
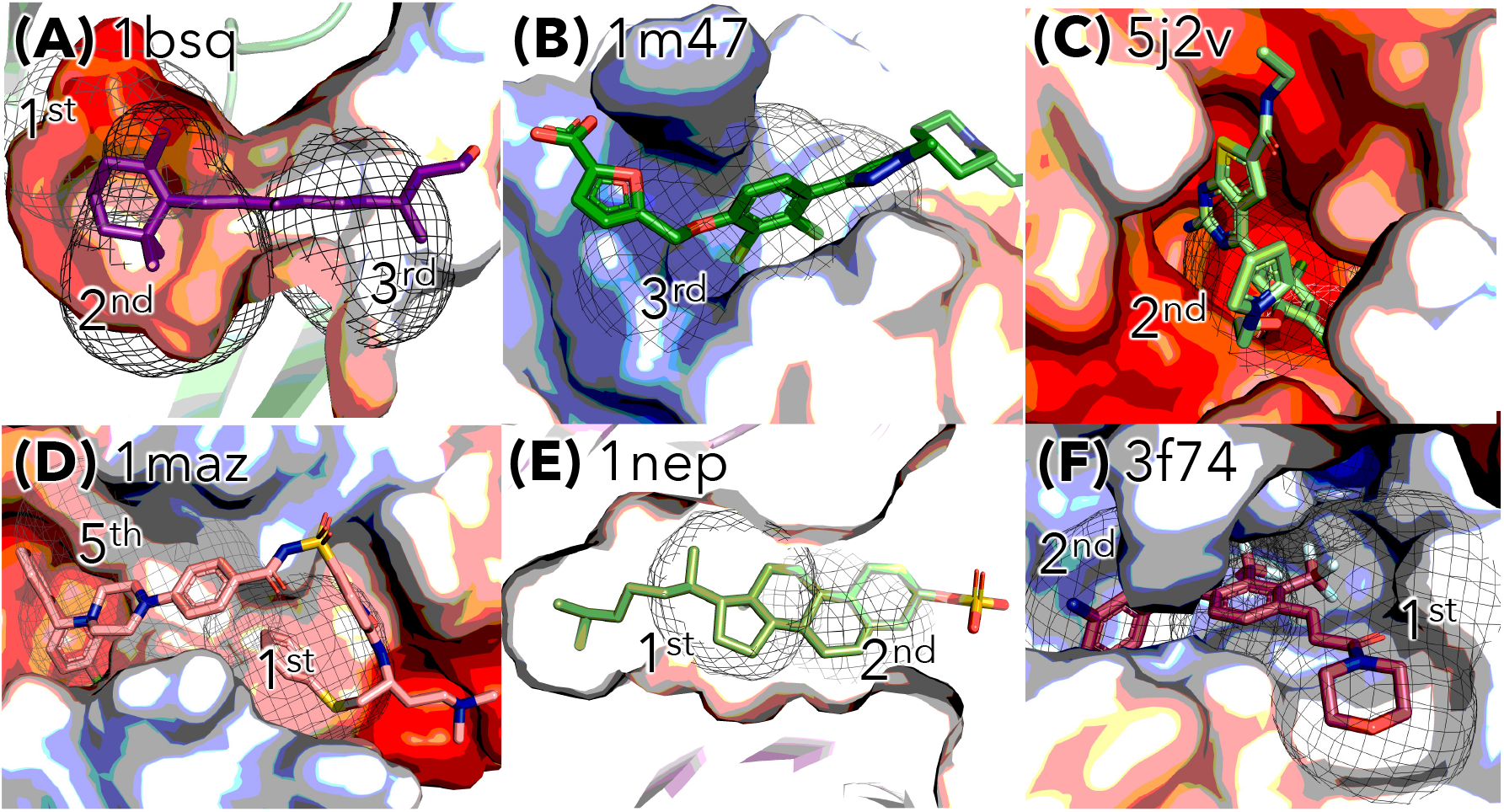
Electrostatic surface of a binding site for each protein. Red regions represent areas of negative electrostatic potential, while blue regions indicate areas of positively electrostatic potential. The meshes show the DBSCAN clusters, corresponding to the clusters of black spheres in Figure 3.

To investigate the interaction preference of xenon with amino-acid residues, I performed contact analysis as follows. (i) The centre-of-mass (COM) point of each residue was calculated. (ii) The distances between COM points and the xenon atoms were measured. (iii) If the distance from the COM point of a residue was less than 4.5 Å, then the xenon atoms were considered to be in contact with the amino-acid residue.

I found that xenon atoms were not only bound to hydrophobic but also to hydrophilic residues via their apolar regions, methylene groups (Figure 5). For example, a xenon atom was caged by residues including Arg (Top of Figure 5B) or Glu and Asp (Bottom of Figure 5B)

**Figure 5:**
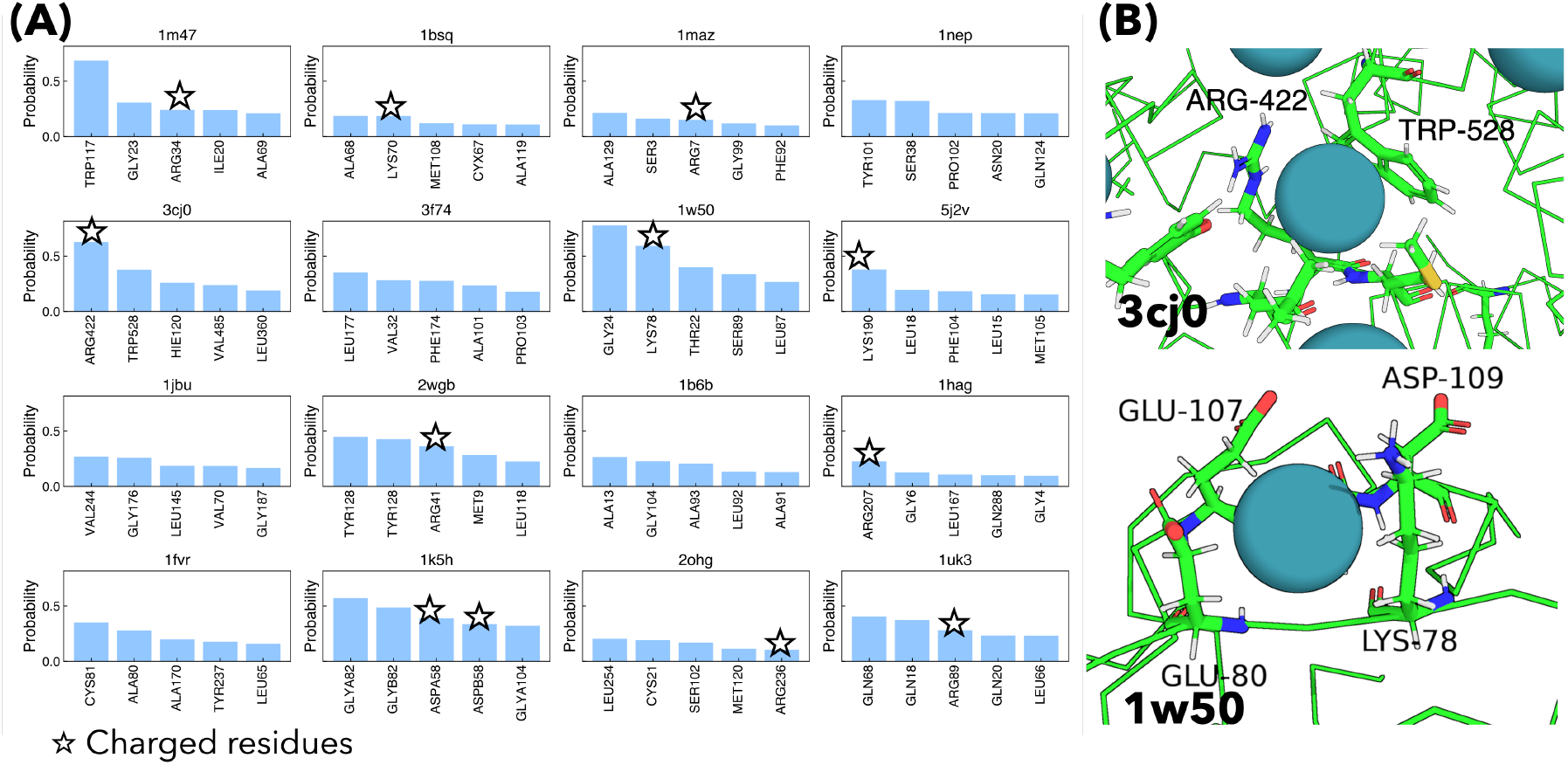
(A) Contact analysis of xenon. The top five frequently-contacting residues are shown in each plot. Each black star indicates hydrophilic residues. (B) Two examples of the way of xenon atoms to interact with hydrophilic residues.

Such methylene-mediated interaction of xenon is also critical for CB site recognition. In fact, two positively-charged residues (R38 and K35) in the PDB ID 1m47 comprise the CB site and the methylene groups caged the xenon cluster (Figure 6).

**Figure 6:**
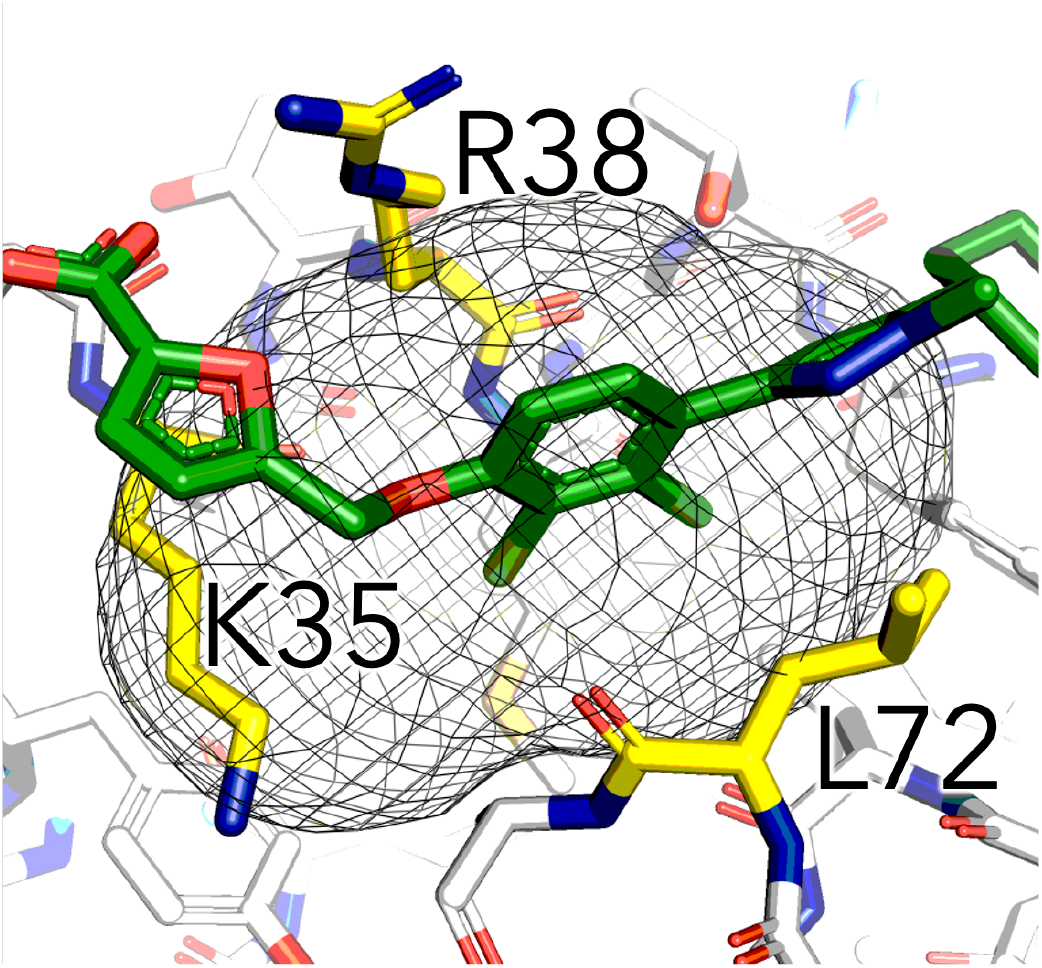
A CB site in the PDBID 1m47. The site includes two charged residues R38 and K35 that create positively-charged surface.

## Limitation

Although xenon is likely to bind to apparent and CB sites, it has limitations as a probe. Firstly, xenon binds not only to CB sites but also to other pockets, leading to the increase in FDR for the identification of CB sites (Tables 4 and 5). This was also observed in the myoglobin system in Figure S8.

To address this, filtering algorithms should be introduced. The top *n* threshold employed in the Tables 4 and 5 would be one that eliminates false positives, yet more sophisticated methods could be developed in the future. In this study, I rather focused on assessing xenon’s intrinsic binding preferences for apparent and CB sites.

Secondly, large conformational changes were not induced by the inclusion of xenon atoms in canonical MD simulations (Figure 7). In Figure 7A, the protein surface is so flat that xenon struggled to settle, and thus the strategy in this work failed to identify the apparent and CB sites. In Figure 7B, the CB site in the apo state is covered with a loop and the loop movement requires movement of a helix, which was not observed in MD simulations. In Figure 7C, the CB site in the apo state is covered with a helix, so that the removal of helix is necessary for the exposure of CB site; however, such large conformational changes were not observed in MD simulations.

**Figure 7:**
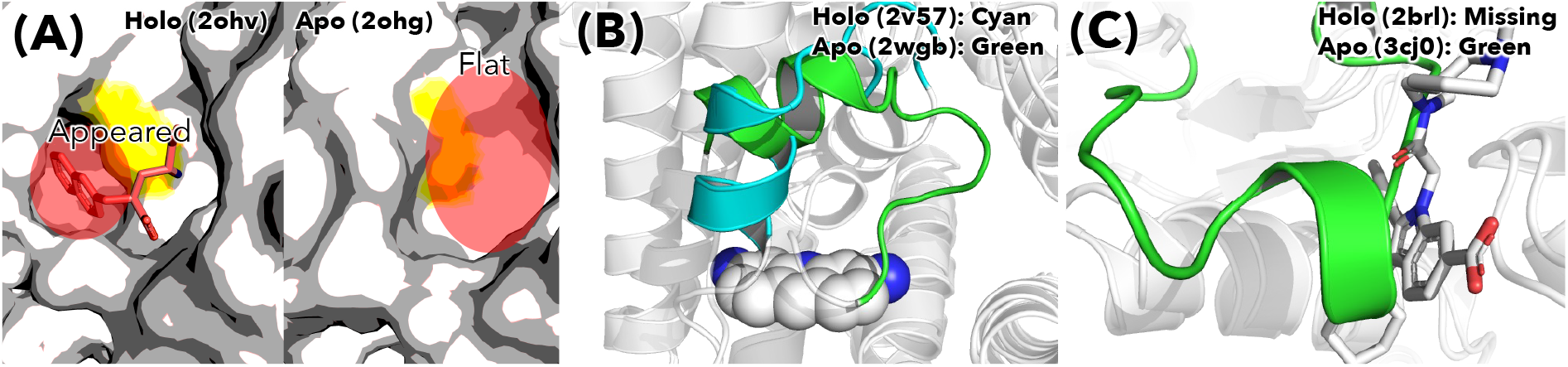
Cases where xenon failed to bind to CB sites. (A): Holo and apo structures are depicted by surface representation and the yellow region indicates amino-acid residues judged as those in a CB site. (B and C): The CB sites in holo and apo structures are coloured by cyan and green, respectively. As shown, the apo structures hides the ligand binding site via a loop or helix conformation. There are missing residues of the holo structure corresponding to the green-coloured helix region (in panel C).

To observe such large conformational changes, the use of an enhanced sampling method would be required for efficient exploration of CB sites via xenon. In fact, Kuzmanic et al. suggested the effectiveness of an enhanced sampling method with cosolvent probes to explore CB sites. ^8^ Although the present study has several limitations, it has the following significance: (i) it provides the evaluation of the xenon model with the LJ parameters for xenon-protein interactions and (ii) it investigates the ability of xenon to bind to apparent binding and CB sites for various proteins.

## Conclusion

In this study, I demonstrated that xenon, through atomistic MD simulations, can probe both apparent and CB sites on proteins. The key finding is that xenon, even in simulations from apo protein structures, is able to identify CB sites if large structural rearrangement is not required. Compared to benzene, xenon achieved higher hit rates in identifying CB sites when more than one cluster was considered, indicating its superior suitability as a probe.

The present work complements the work by Vithani et al. who utilised xenon as a probe and performed MD simulations for a protein KRAS to identify its CB sites. The authors state that “we believe xenon might make a reasonable generic probe of cryptic sites within any protein complex, including KRAS.”. ^19^ While their work was a detailed case study on a single protein, my work solidified their statement and assessed more generic applicability of xenon across sixteen different proteins with known CB sites.

The results in this study suggest that xenon is not only a viable but a potentially superior probe for detecting CB sites in atomistic MD simulations. Its effectiveness, even from apo structures, highlights its utility in scenarios where ligand-bound information is unavailable. Future work could involve leveraging curated datasets such as CryptoBench ^79^ for more systematic benchmarking. Moreover, it is crucial to compare computational results to those of experiments: A plausible collaboration would involve the combination between MD and NMR-based strategy, which can identify the population of CB sites in equilibrium. ^80^

Taken together, this study reinforces the role of xenon as a powerful tool for CB site identification and provides a foundation for its broader application in computational drug discovery.

## Associated Content

### Data and Software Availability

The Gromacs topology file of xenon is publicly available at https://gist.github.com/physicshinzui/718b101964d44a1d58dfd4cc8bd15860. All the MD simulation data (xtc format trajectory files and their reference PDBs) are also publicly available at https://zenodo.org/records/17299983. Due to the data size, all the water molecules in each MD data were removed.

### Supporting Information

The Supporting Information is available free of charge. Supporting Information: Five trajectories of RMSD for each target proteins with xenons (Figure S1); Five trajectories of RMSD for each target proteins with benzenes (Figure S2); HDBSCAN parameter search for xenon (Figure S3); HDBSCAN parameter search for benzene (Figure S4); Free-energy iso-surface of xenon (Figure S5); Five trajectories of RMSD for each target proteins with xenon (Figure S6); Success and failed cases of CB sites identification by benzene (Figure S7); Overall picture of free-energy iso-surface of xenon in the myoglobin system (Figure S8); Table S1-S7

### Author Contributions

The author confirms sole responsibility for the following: study conception and design, data collection, analysis and interpretation of results, and manuscript preparation.

## Acknowledgement

This work was supported by Kitasato University Academic Research Encouragement Fund. The computation was performed using the Research Center for Computational Science, Okazaki, Japan (Project: 24-IMS-C128). I would like to thank Dr. Yoshifumi Fukunishi for fruitful discussions. I would like to thank Dr. Yasuhiro Imada for his help in generating the topology for the myoglobin system.

## References

(1) Le Guilloux, V.; Schmidtke, P.; Tuffery, P. Fpocket: an open source platform for ligand pocket detection. BMC Bioinformatics 2009, 10, 168.

(2) Hendlich, M.; Rippmann, F.; Barnickel, G. LIGSITE: automatic and efficient detection of potential small molecule-binding sites in proteins. J. Mol. Graph. Model. 1997, 15, 359–63, 389.

(3) Fukunishi, Y.; Nakamura, H. Prediction of ligand-binding sites of proteins by molecular docking calculation for a random ligand library: Prediction of Ligand-Binding Sites. Protein Sci. 2011, 20, 95–106.

(4) Frembgen-Kesner, T.; Elcock, A. H. Computational sampling of a cryptic drug binding site in a protein receptor: explicit solvent molecular dynamics and inhibitor docking to p38 MAP kinase. J. Mol. Biol. 2006, 359, 202–214.

(5) Cimermancic, P.; Weinkam, P.; Rettenmaier, T. J.; Bichmann, L.; Keedy, D. A.; Woldeyes, R. A.; Schneidman-Duhovny, D.; Demerdash, O. N.; Mitchell, J. C.; Wells, J. A.; Fraser, J. S.; Sali, A. CryptoSite: Expanding the druggable proteome by characterization and prediction of cryptic binding sites. J. Mol. Biol. 2016, 428, 709–719.

(6) Meller, A.; Ward, M.; Borowsky, J.; Kshirsagar, M.; Lotthammer, J. M.; Oviedo, F.; Ferres, J. L.; Bowman, G. R. Predicting locations of cryptic pockets from single protein structures using the PocketMiner graph neural network. Nat. Commun. 2023, 14, 1177.

(7) Škrhák, V.; Riedlova, K.; Novotný, M.; Hoksza, D. Cryptic binding site prediction with protein language models. 2023 IEEE International Conference on Bioinformatics and Biomedicine (BIBM). 2023; pp 2883–2888.

(8) Kuzmanic, A.; Bowman, G. R.; Juarez-Jimenez, J.; Michel, J.; Gervasio, F. L. Investigating cryptic binding sites by molecular dynamics simulations. Acc. Chem. Res. 2020, 53, 654–661.

(9) Iida, S.; Nakamura, H. K.; Mashimo, T.; Fukunishi, Y. Structural fluctuations of aromatic residues in an APO-form reveal cryptic binding sites: Implications for fragment-based drug design. J. Phys. Chem. B 2020, 124, 9977–9986.

(10) Meller, A.; Bhakat, S.; Solieva, S.; Bowman, G. R. Accelerating cryptic pocket discovery using AlphaFold. J. Chem. Theory Comput. 2023, 19, 4355–4363.

(11) Yanagisawa, K.; Moriwaki, Y.; Terada, T.; Shimizu, K. EXPRORER: Rational cosolvent set construction method for cosolvent molecular dynamics using large-scale computation. J. Chem. Inf. Model. 2021, 61, 2744–2753.

(12) Smith, R. D.; Carlson, H. A. Identification of cryptic binding sites using MixMD with standard and accelerated molecular dynamics. J. Chem. Inf. Model. 2021, 61, 1287– 1299.

(13) Koseki, J.; Motono, C.; Yanagisawa, K.; Kudo, G.; Yoshino, R.; Hirokawa, T.; Imai, K. CrypToth: Cryptic pocket detection through mixed-solvent molecular dynamics simulations based topological data analysis. bioRxiv 2024,

(14) Motono, C.; Yanagisawa, K.; Koseki, J.; Imai, K. CrypTothML: An integrated mixed-solvent molecular dynamics simulation and machine learning approach for cryptic site prediction. Int. J. Mol. Sci. 2025, 26.

(15) Tan, Y. S.; Reeks, J.; Brown, C. J.; Thean, D.; Ferrer Gago, F. J.; Yuen, T. Y.; Goh, E. T. L.; Lee, X. E. C.; Jennings, C. E.; Joseph, T. L.; Lakshminarayanan, R.; Lane, D. P.; Noble, M. E. M.; Verma, C. S. Benzene probes in molecular dynamics simulations reveal novel binding sites for ligand design. J. Phys. Chem. Lett. 2016, 7, 3452–3457.

(16) Zuzic, L.; Marzinek, J. K.; Warwicker, J.; Bond, P. J. A benzene-mapping approach for uncovering cryptic pockets in membrane-bound proteins. J. Chem. Theory Comput. 2020, 16, 5948–5959.

(17) Martinez-Rosell, G.; Lovera, S.; Sands, Z. A.; De Fabritiis, G. PlayMolecule Cryptic-Scout: Predicting protein cryptic sites using mixed-solvent molecular simulations. J. Chem. Inf. Model. 2020, 60, 2314–2324.

(18) Prangé, T.; Schiltz, M.; Pernot, L.; Colloc’h, N.; Longhi, S.; Bourguet, W.; Fourme, R. Exploring hydrophobic sites in proteins with xenon or krypton. Proteins 1998, 30, 61–73.

(19) Vithani, N.; Zhang, S.; Thompson, J. P.; Patel, L. A.; Demidov, A.; Xia, J.; Balaeff, A.; Mentes, A.; Arnautova, Y. A.; Kohlmann, A.; Lawson, J. D.; Nicholls, A.; Skillman, A. G.; LeBard, D. N. Exploration of cryptic pockets using enhanced sampling along normal modes: A case study of KRAS G12D. J. Chem. Inf. Model. 2024, 64, 8258–8273.

(20) Roose, B. W.; Zemerov, S. D.; Dmochowski, I. J. Xenon-protein interactions: Characterization by X-ray crystallography and hyper-CEST NMR. Methods Enzymol. 2018, 602, 249–272.

(21) Wiebelhaus, N.; Singh, N.; Zhang, P.; Craig, S. L.; Beratan, D. N.; Fitzgerald, M. C. Discovery of the xenon-protein interactome using large-scale measurements of protein folding and stability. J. Am. Chem. Soc. 2022, 144, 3925–3938.

(22) Harris, P. D.; Barnes, R. The uses of helium and xenon in current clinical practice. Anaesthesia 2008, 63, 284–293.

(23) Dickinson, R.; Franks, N. P. Bench-to-bedside review: Molecular pharmacology and clinical use of inert gases in anesthesia and neuroprotection. Crit. Care 2010, 14, 229.

(24) Tilton, R. F., Jr; Singh, U. C.; Weiner, S. J.; Connolly, M. L.; Kuntz, I. D., Jr; Kollman, P. A.; Max, N.; Case, D. A. Computational studies of the interaction of myoglobin and xenon. J. Mol. Biol. 1986, 192, 443–456.

(25) Mann, G.; Hermans, J. Modeling protein-small molecule interactions: structure and thermodynamics of noble gases binding in a cavity in mutant phage T4 lysozyme L99A. J. Mol. Biol. 2000, 302, 979–989.

(26) Rutkai, G.; Thol, M.; Span, R.; Vrabec, J. How well does the Lennard-Jones potential represent the thermodynamic properties of noble gases? Mol. Phys. 2017, 115, 1104– 1121.

(27) GROMACS 2022.4 manual; nonbonded-interactions. https://manual.gromacs.org/2022.4/reference-manual/functions/nonbonded-interactions.html, Accessed: 2023-12-9.

(28) Tang, K. T.; Toennies, J. P. New combining rules for well parameters and shapes of the van der Waals potential of mixed rare gas systems. Z. Phys. D At. Mol. Clust. 1986, 1, 91–101.

(29) Roose, B. W.; Zemerov, S. D.; Wang, Y.; Kasimova, M. A.; Carnevale, V.; Dmochowski, I. J. A structural basis for 129 Xe hyper-CEST signal in TEM-1 β-lactamase. Chemphyschem 2019, 20, 260–267.

(30) Abraini, J. H.; Marassio, G.; David, H. N.; Vallone, B.; Prangé, T.; Colloc’h, N. Crystallographic studies with xenon and nitrous oxide provide evidence for protein-dependent processes in the mechanisms of general anesthesia. Anesthesiology 2014, 121, 1018– 1027.

(31) Hickman, A. B.; Klein, D. C.; Dyda, F. Melatonin biosynthesis: the structure of serotonin N-acetyltransferase at 2.5 A resolution suggests a catalytic mechanism. Mol. Cell 1999, 3, 23–32.

(32) Wolf, E.; De Angelis, J.; Khalil, E. M.; Cole, P. A.; Burley, S. K. X-ray crystallographic studies of serotonin N-acetyltransferase catalysis and inhibition. J. Mol. Biol. 2002, 317, 215–224.

(33) Qin, B. Y.; Bewley, M. C.; Creamer, L. K.; Baker, E. N.; Jameson, G. B. Functional implications of structural differences between variants A and B of bovine beta-lactoglobulin. Protein Sci. 1999, 8, 75–83.

(34) Kontopidis, G.; Holt, C.; Sawyer, L. The ligand-binding site of bovine beta-lactoglobulin: evidence for a function? J. Mol. Biol. 2002, 318, 1043–1055.

(35) Shewchuk, L. M.; Hassell, A. M.; Ellis, B.; Holmes, W. D.; Davis, R.; Horne, E. L.; Kadwell, S. H.; McKee, D. D.; Moore, J. T. Structure of the Tie2 RTK domain: self-inhibition by the nucleotide binding loop, activation loop, and C-terminal tail. Structure 2000, 8, 1105–1113.

(36) Hodous, B. L. et al. Synthesis, structural analysis, and SAR studies of triazine derivatives as potent, selective Tie-2 inhibitors. Bioorg. Med. Chem. Lett. 2007, 17, 2886– 2889.

(37) Antonysamy, S. S. et al. Fragment-based discovery of hepatitis C virus NS5b RNA polymerase inhibitors. Bioorg. Med. Chem. Lett. 2008, 18, 2990–2995.

(38) Di Marco, S.; Volpari, C.; Tomei, L.; Altamura, S.; Harper, S.; Narjes, F.; Koch, U.; Rowley, M.; De Francesco, R.; Migliaccio, G.; Carfí, A. Interdomain communication in hepatitis C virus polymerase abolished by small molecule inhibitors bound to a novel allosteric site. J. Biol. Chem. 2005, 280, 29765–29770.

(39) Vijayalakshmi, J.; Padmanabhan, K. P.; Mann, K. G.; Tulinsky, A. The isomorphous structures of prethrombin2, hirugen-, and PPACK-thrombin: changes accompanying activation and exosite binding to thrombin. Protein Sci. 1994, 3, 2254–2271.

(40) Eigenbrot, C.; Kirchhofer, D.; Dennis, M. S.; Santell, L.; Lazarus, R. A.; Stamos, J.; Ultsch, M. H. The factor VII zymogen structure reveals reregistration of beta strands during activation. Structure 2001, 9, 627–636.

(41) Kadono, S. et al. Structure-based design of P3 moieties in the peptide mimetic factor VIIa inhibitor. Biochem. Biophys. Res. Commun. 2005, 327, 589–596.

(42) Reuter, K.; Sanderbrand, S.; Jomaa, H.; Wiesner, J.; Steinbrecher, I.; Beck, E.; Hintz, M.; Klebe, G.; Stubbs, M. T. Crystal structure of 1-deoxy-D-xylulose-5-phosphate reductoisomerase, a crucial enzyme in the non-mevalonate pathway of isoprenoid biosynthesis. J. Biol. Chem. 2002, 277, 5378–5384.

(43) Yajima, S.; Hara, K.; Iino, D.; Sasaki, Y.; Kuzuyama, T.; Ohsawa, K.; Seto, H. Structure of 1-deoxy-D-xylulose 5-phosphate reductoisomerase in a quaternary complex with a magnesium ion, NADPH and the antimalarial drug fosmidomycin. Acta Crystallogr. Sect. F Struct. Biol. Cryst. Commun. 2007, 63, 466–470.

(44) Kim, K.-H.; Bong, Y.-J.; Park, J. K.; Shin, K.-J.; Hwang, K. Y.; Kim, E. E. Structural basis for glutamate racemase inhibition. J. Mol. Biol. 2007, 372, 434–443.

(45) Arkin, M. R.; Randal, M.; DeLano, W. L.; Hyde, J.; Luong, T. N.; Oslob, J. D.; Raphael, D. R.; Taylor, L.; Wang, J.; McDowell, R. S.; Wells, J. A.; Braisted, A. C. Binding of small molecules to an adaptive protein-protein interface. Proc. Natl. Acad. Sci. U. S. A. 2003, 100, 1603–1608.

(46) Thanos, C. D.; Randal, M.; Wells, J. A. Potent small-molecule binding to a dynamic hot spot on IL-2. J. Am. Chem. Soc. 2003, 125, 15280–15281.

(47) Muchmore, S. W.; Sattler, M.; Liang, H.; Meadows, R. P.; Harlan, J. E.; Yoon, H. S.; Nettesheim, D.; Chang, B. S.; Thompson, C. B.; Wong, S. L.; Ng, S. L.; Fesik, S. W. X-ray and NMR structure of human Bcl-xL, an inhibitor of programmed cell death. Nature 1996, 381, 335–341.

(48) Lee, E. F.; Czabotar, P. E.; Smith, B. J.; Deshayes, K.; Zobel, K.; Colman, P. M.; Fairlie, W. D. Crystal structure of ABT-737 complexed with Bcl-xL: implications for selectivity of antagonists of the Bcl-2 family. Cell Death Differ. 2007, 14, 1711–1713.

(49) Friedland, N.; Liou, H.-L.; Lobel, P.; Stock, A. M. Structure of a cholesterol-binding protein deficient in Niemann-Pick type C2 disease. Proc. Natl. Acad. Sci. U. S. A. 2003, 100, 2512–2517.

(50) Zhang, H.; Astrof, N. S.; Liu, J.-H.; Wang, J.-H.; Shimaoka, M. Crystal structure of isoflurane bound to integrin LFA-1 supports a unified mechanism of volatile anesthetic action in the immune and central nervous systems. FASEB J. 2009, 23, 2735–2740.

(51) Guckian, K. M.; Lin, E. Y.-S.; Silvian, L.; Friedman, J. E.; Chin, D.; Scott, D. M. Design and synthesis of a series of meta aniline-based LFA-1 ICAM inhibitors. Bioorg. Med. Chem. Lett. 2008, 18, 5249–5251.

(52) Yang, H.; Yang, M.; Ding, Y.; Liu, Y.; Lou, Z.; Zhou, Z.; Sun, L.; Mo, L.; Ye, S.; Pang, H.; Gao, G. F.; Anand, K.; Bartlam, M.; Hilgenfeld, R.; Rao, Z. The crystal structures of severe acute respiratory syndrome virus main protease and its complex with an inhibitor. Proc. Natl. Acad. Sci. U. S. A. 2003, 100, 13190–13195.

(53) Lu, I.-L.; Mahindroo, N.; Liang, P.-H.; Peng, Y.-H.; Kuo, C.-J.; Tsai, K.-C.; Hsieh, H.-P.; Chao, Y.-S.; Wu, S.-Y. Structure-based drug design and structural biology study of novel nonpeptide inhibitors of severe acute respiratory syndrome coronavirus main protease. J. Med. Chem. 2006, 49, 5154–5161.

(54) Patel, S.; Vuillard, L.; Cleasby, A.; Murray, C. W.; Yon, J. Apo and inhibitor complex structures of BACE (beta-secretase). J. Mol. Biol. 2004, 343, 407–416.

(55) Björklund, C.; Oscarson, S.; Benkestock, K.; Borkakoti, N.; Jansson, K.; Lindberg, J.; Vrang, L.; Hallberg, A.; Rosenquist, A.; Samuelsson, B. Design and synthesis of potent and selective BACE-1 inhibitors. J. Med. Chem. 2010, 53, 1458–1464.

(56) Bellinzoni, M.; Buroni, S.; Schaeffer, F.; Riccardi, G.; De Rossi, E.; Alzari, P. M. Structural plasticity and distinct drug-binding modes of LfrR, a mycobacterial efflux pump regulator. J. Bacteriol. 2009, 191, 7531–7537.

(57) Amaral, M.; Kokh, D. B.; Bomke, J.; Wegener, A.; Buchstaller, H. P.; Eggenweiler, H. M.; Matias, P.; Sirrenberg, C.; Wade, R. C.; Frech, M. Protein conformational flexibility modulates kinetics and thermodynamics of drug binding. Nat. Commun. 2017, 8.

(58) Brough, P. A. et al. Combining hit identification strategies: fragment-based and in silico approaches to orally active 2-aminothieno[2,3-d]pyrimidine inhibitors of the Hsp90 molecular chaperone. J. Med. Chem. 2009, 52, 4794–4809. modules/pymol/util.py at 03d7a7fcf0bd95cd93d710a1268dbace2ed77765 · schrodinger/pymol-open-source.

(59) modules/pymol/util.py at 03d7a7fcf0bd95cd93d710a1268dbace2ed77765. schrodinger/pymol-open-source.

(60) Shirts, M. R.; Klein, C.; Swails, J. M.; Yin, J.; Gilson, M. K.; Mobley, D. L.; Case, D. A.; Zhong, E. D. Lessons learned from comparing molecular dynamics engines on the SAMPL5 dataset. J. Comput. Aided Mol. Des. 2017, 31, 147–161.

(61) Páll, S.; Hess, B. A flexible algorithm for calculating pair interactions on SIMD architectures. arXiv [physics.comp-ph] 2013,

(62) Hess, B.; Bekker, H.; Berendsen, H. J. C.; Fraaije, J. G. E. M. LINCS: A linear constraint solver for molecular simulations. J. Comput. Chem. 1997, 18, 1463–1472.

(63) Bussi, G.; Donadio, D.; Parrinello, M. Canonical sampling through velocity rescaling. J. Chem. Phys. 2007, 126, 014101.

(64) Berendsen, H. J. C.; Postma, J. P. M.; van Gunsteren, W. F.; DiNola, A.; Haak, J. R. Molecular dynamics with coupling to an external bath. J. Chem. Phys. 1984, 81, 3684– 3690.

(65) Maier, J. A.; Martinez, C.; Kasavajhala, K.; Wickstrom, L.; Hauser, K. E.; Simmerling, C. Ff14SB: Improving the accuracy of protein side chain and backbone parameters from ff99SB. J. Chem. Theory Comput. 2015, 11, 3696–3713.

(66) Jorgensen, W. L.; Chandrasekhar, J.; Madura, J. D.; Impey, R. W.; Klein, M. L. Comparison of simple potential functions for simulating liquid water. J. Chem. Phys. 1983, 79, 926–935.

(67) Essmann, U.; Perera, L.; Berkowitz, M. L.; Darden, T.; Lee, H.; Pedersen, L. G. A smooth particle mesh Ewald method. J. Chem. Phys. 1995, 103, 8577–8593.

(68) Bauer, P.; Hess, B.; Lindahl, E. GROMACS 2022.4 Manual. 2022.

(69) Sousa da Silva, A.W.; Vranken, W. F. ACPYPE -AnteChamber PYthon Parser interfacE. BMC Res. Notes 2012, 5, 367.

(70) Anandakrishnan, R.; Aguilar, B.; Onufriev, A. V. H++ 3.0: automating pK prediction and the preparation of biomolecular structures for atomistic molecular modeling and simulations. Nucleic Acids Res. 2012, 40, W537–41.

(71) Li, P.; Merz, K. M., Jr MCPB.Py: A python based metal center parameter builder. J. Chem. Inf. Model. 2016, 56, 599–604.

(72) Case, D. A. et al. AmberTools. J. Chem. Inf. Model. 2023, 63, 6183–6191.

(73) Frisch, M. J. et al. Gaussian 16 ; Gaussian, Inc: Wallingford CT, 2016.

(74) Becke, A. D. Density-functional thermochemistry. III. The role of exact exchange. J. Chem. Phys. 1993, 98, 5648–5652.

(75) Campello, R. J. G. B.; Moulavi, D.; Sander, J. Advances in Knowledge Discovery and Data Mining ; Lecture notes in computer science; Springer Berlin Heidelberg: Berlin, Heidelberg, 2013; pp 160–172.

(76) Michaud-Agrawal, N.; Denning, E. J.; Woolf, T. B.; Beckstein, O. MDAnalysis: a toolkit for the analysis of molecular dynamics simulations. J. Comput. Chem. 2011, 32, 2319–2327.

(77) Gowers, R.; Linke, M.; Barnoud, J.; Reddy, T.; Melo, M.; Seyler, S.; Domański, J.; Dotson, D.; Buchoux, S.; Kenney, I.; Beckstein, O. MDAnalysis: A python package for the rapid analysis of molecular dynamics simulations. Proceedings of the Python in Science Conference. 2016; pp 98–105.

(78) Jurrus, E. et al. Improvements to the APBS biomolecular solvation software suite. Protein Sci. 2018, 27, 112–128.

(79) Škrhák, V.; Novotný, M.; Feidakis, C. P.; Krivák, R.; Hoksza, D. CryptoBench: cryptic protein-ligand binding sites dataset and benchmark. Bioinformatics 2024, 41.

(80) Mizukoshi, Y.; Takeuchi, K.; Tokunaga, Y.; Matsuo, H.; Imai, M.; Fujisaki, M.; Kamoshida, H.; Takizawa, T.; Hanzawa, H.; Shimada, I. Targeting the cryptic sites: NMR-based strategy to improve protein druggability by controlling the conformational equilibrium. Sci. Adv. 2020, 6.

