## Supporting Information for "Xenon Explores Apparent and Cryptic Binding Sites"

Shinji Iida[a*]

a. *Department of Data Science, Kitasato University, Kanagawa, 252-0373, Japan*

#### List of Figures

S1 Five trajectories of RMSD for each target proteins with xenons . . . . . . . . S3
S2 Five trajectories of RMSD for each target proteins with benzenes . . . . . . S4
S3 HDBSCAN parameter search for xenon . . . . . . . . . . . . . . . . . . . . . . S7
S4 HDBSCAN parameter search for benzene . . . . . . . . . . . . . . . . . . . . . S8
S5 Free-energy iso-surface of xenon . . . . . . . . . . . . . . . . . . . . . . . . . S10
S6 Five trajectories of RMSD for each target proteins with xenon . . . . . . . . S11
S7 Success and failed cases of CB sites identification by benzene molecule . . . . S12
S8 Overall picture of free-energy iso-surface of xenon in the myoglobin system . S13

#### List of Tables

S1 Set of amino-acid residues in CB sites . . . . . . . . . . . . . . . . . . . . . . S5
S2 Set of amino-acid residues in apparent binding sites . . . . . . . . . . . . . . . S6

S3 Parameters of HDBSCAN . . . . . . . . . . . . . . . . . . . . . . . . . . . . S8
S4 Identification of **apparent binding sites** by **xenons** depending on the top $n$ clusters . . . . . . . . . . . . . . . . . . . . . . . . . . . . . . . . . S14
S5 Identification of **apparent binding sites** by **benzenes** depending on the top $n$ clusters . . . . . . . . . . . . . . . . . . . . . . . . . . . . . . . S15
S6 Identification of **cryptic binding sites** by **xenons** depending on the top $n$ clusters . . . . . . . . . . . . . . . . . . . . . . . . . . . . . . . . . . S15
S7 Identification of **cryptic binding sites** by **benzenes** depending on the top $n$ clusters . . . . . . . . . . . . . . . . . . . . . . . . . . . . . . . . . S16

### Root mean squared deviation during MD simulations

The insertion of xenons/benzenes may cause an artefact during MD simulations, such as unexpected denaturation. To inspect this, I examined if the concentration of the xenon and benzene is appropriate by root-mean squared deviation (RMSD) of each MD snapshots of each target system in Figure S1 and S2. The RMSD was calculated with respect to the backbone atoms (*gmx rms* command was used for this calculation.). Figure S1 and S2 show that the whole structure of each protein was kept during MD simulations.

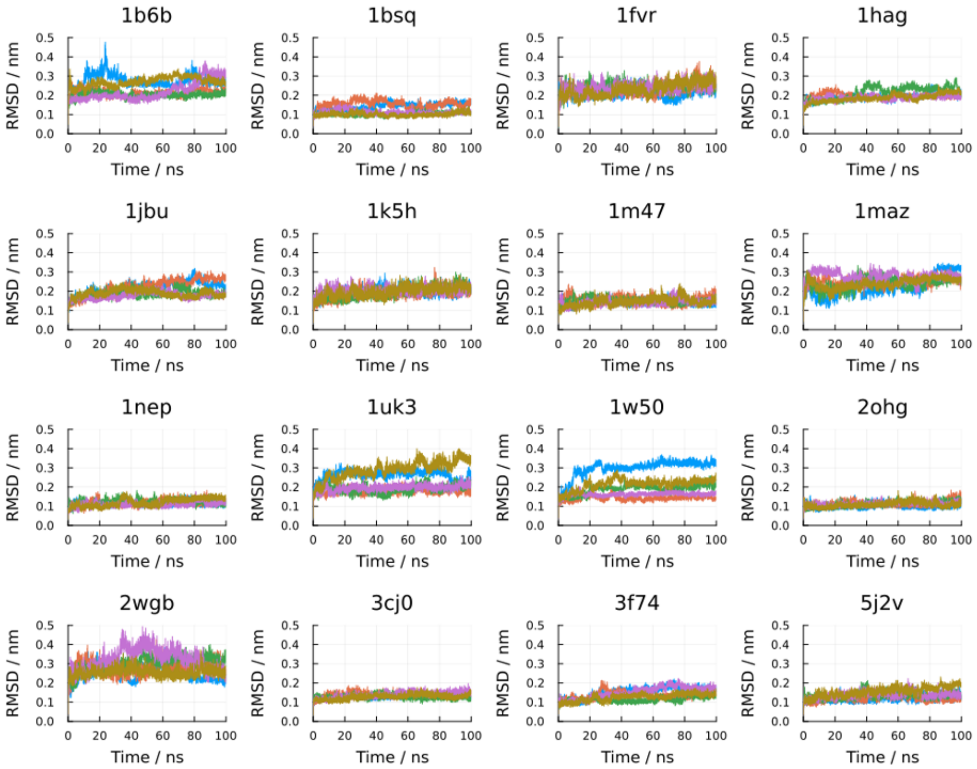


Figure S1: Five trajectories of RMSD for each target proteins with xenons at 300 K. The header in each plot indicates the PDB ID of the apo state.

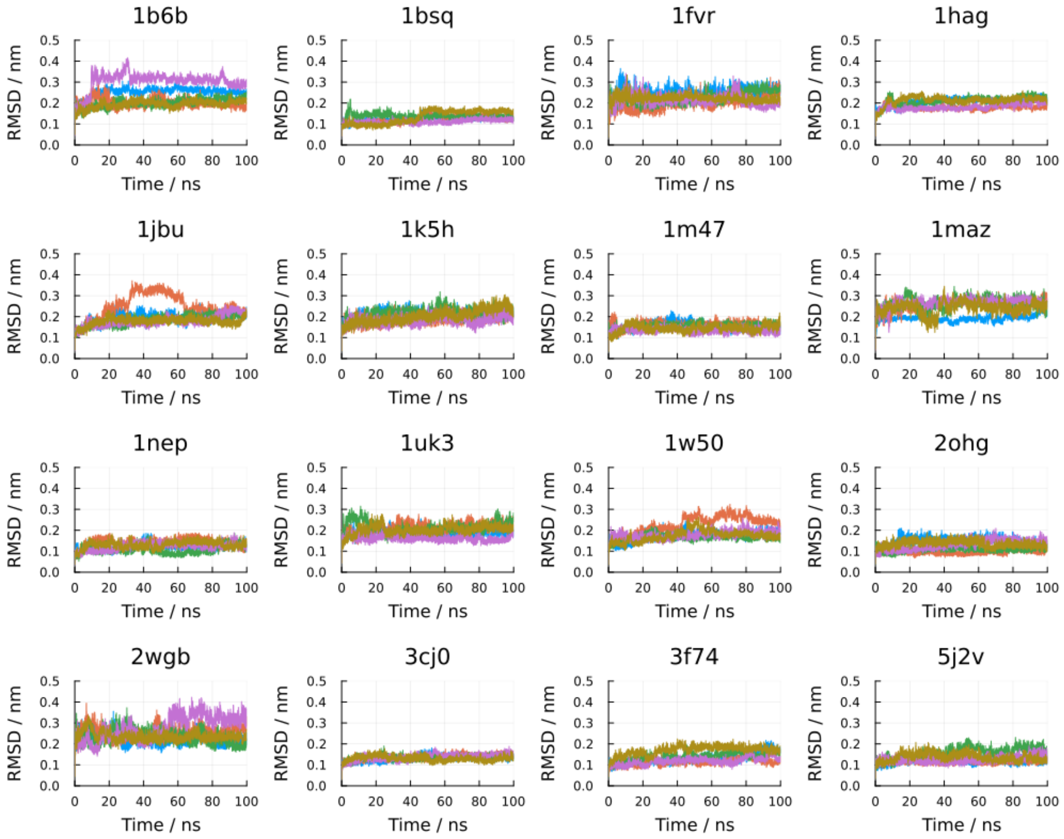


Figure S2: Five trajectories of RMSD for each target proteins with benzenes at 300 K. The header in each plot indicates the PDB ID of an apo state.

Table S1: Set of amino-acid residues in CB sites

| PDBID (apo_holo) | $S_{\mathrm{CB}}$ | | | | | | | |
|---|---|---|---|---|---|---|---|---|
| 1b6b_1kuv | PHE56 | ASN62 | PRO64 | ALA125 | GLY136 | SER137 | GLU161 | ALA163 |
| 1hag_1ghy | TYR60 | TRP60 | GLU192 | GLY216 | GLY219 | | | |
| 1m47_1py2 | LYS35 | ARG38 | TYR45 | LEU72 | | | | |
| 1uk3_2gz7 | HIS41 | | | | | | | |
| 1bsq_1gx8 | LEU39 | MET107 | | | | | | |
| 1jbu_1wun | THR98 | CYS191 | LYS192 | GLY216 | GLN217 | GLY219 | CYS220 | |
| 1maz_2yxj | ALA93 | PHE97 | PHE105 | VAL126 | GLU129 | LEU130 | GLY138 | TYR195 |
| 1w50_3ixj | GLY59 | GLY61 | THR280 | | | | | |
| 1fvr_2oo8 | ALA853 | ASP982 | PHE983 | | | | | |
| 1k5h_2egh | GLY10 | SER11 | GLY35 | LYS36 | VAL101 | SER185 | | |
| 1nep_2hka | GLY57 | PHE66 | LEU94 | TYR100 | | | | |
| 2wgb_2v57 | HIS67 | ASP125 | TRP152 | GLN156 | | | | |
| 3cj0_2brl | LEU492 | | | | | | | |
| 2ohg_2ohv | SER11 | | | | | | | |
| 3f74_3bqm | TYR257 | LYS287 | LEU302 | | | | | |
| 5j2v_2wi7 | PHE138 | | | | | | | |

Table S2: Set of amino-acid residues in apparent binding sites

| PDBID (apo_holo) | $S_{AB}$ |
|---|---|
| 1b6b_1kuv | ALA55, SER60, CYS63, ALA123, LEU124, VAL126, ARG131, GLN132, GLY134, LYS135, MET159, CYS160, PHE167 |
| 1hag_1ghy | ASP189, ALA190, CYS191, ASP194, SER195, CYS220, GLY226 |
| 1m47_1py2 | PRO34, MET39, THR41, PHE42, LYS43, PHE44, GLU62, PRO65, VAL69, ALA73 |
| 1uk3_2gz7 | PRO39, CYS145, HIS164, MET165, GLU166, LEU167, ASP187, GLN189, THR190, GLN192 |
| 1bsq_1gx8 | PHE105, ALA118, GLN120 |
| 1jbu_1wun | HIS57, GLY97, SER170, ASP189, SER190, SER214, TRP215, GLY226 |
| 1maz_2yxj | GLU96, TYR101, ALA104, ASN136, ARG139, VAL141, ALA142, SER145 |
| 1w50_3ixj | ASP80, GLY82, SER83, PRO118, TYR119, THR120, GLN121, GLY122, PHE156, ILE174, ASP276, GLY278, THR279, ASN281, ARG283 |
| 1fvr_2oo8 | LYS855, GLU872, ILE886, ILE902, TYR904, ALA905, HIS962, ILE980, ALA981, PHE983 |
| 1k5h_2egh | GLY7, SER8, THR9, ILE12, ALA34, ASN37, ASP56, ALA99, ILE100, ALA104, ALA122, ASN123, LYS124, ASP149, SER150, GLU151, GLY184, TRP211, SER221, MET275 |
| 1nep_2hka | VAL64 |
| 2wgb_2v57 | SER70, ASN71, TYR106, GLY124, ARG148, MET155 |
| 3cj0_2brl | LEU392, ALA393, ALA395, ALA396, LEU425, HIS428, GLY493, VAL494 |
| 2ohg_2ohv | GLY12, ASP36, ARG39, ILE50, TYR53, THR54, THR75, ALA76, THR117, MET119 |
| 3f74_3bqm | VAL157, VAL286 |
| 5j2v_2wi7 | ASN51, SER52, ALA55, ILE96, GLY97, MET98, GLY135, THR184 |

#### Parameter search of HDBSCAN

HDBSCAN has the primary parameter, minimum cluster size *min_clust_size*, which, in our context, sets the minimum number of atoms in a cluster in the Euclidean space. Since each simulation system did not include the same number of xenons/benzenes, it is necessary to normalise the minimum cluster size to employ coherent value of *min_clust_size* for each system. For this reason, I defined *min_clust_size* by $wN_{\text{mol}}$ where, for each system, $N_{\text{mol}}$ is the number of xenons/benzenes and $w$ is a weight. The weight was grid-searched, and it was found that 0.0025 was optimal based on the criteria of elbow method (Figure S3 and S4). The other parameters of HDBSCAN were set to the default values (Table S3).

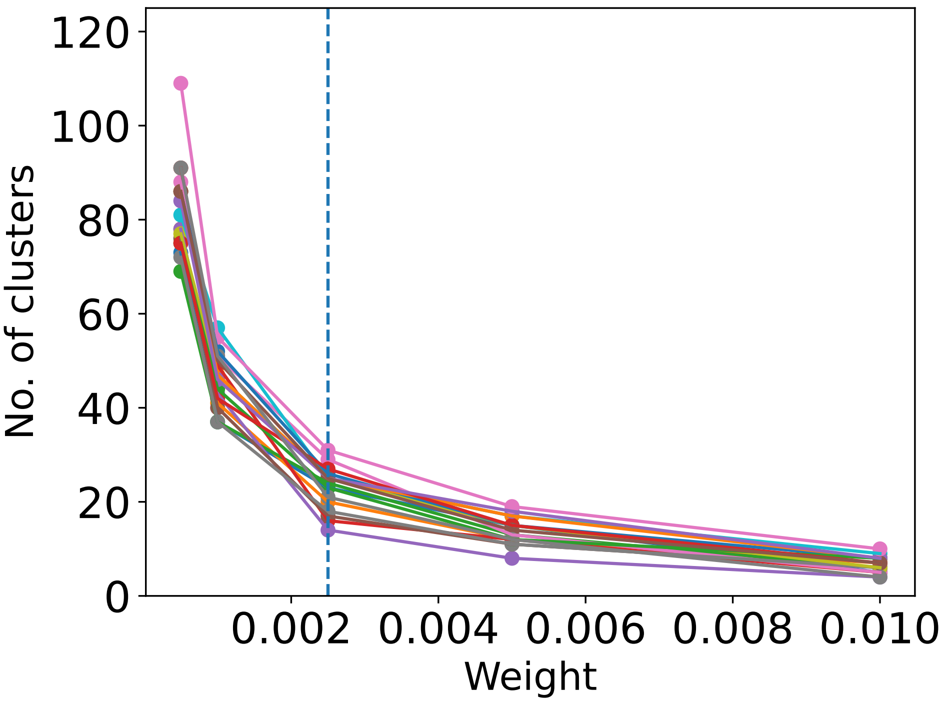


Figure S3: Parameter search for *min_clust_size*. Each coloured line indicates the dependency on weight for a simulation system. The dashed line points to the optimal value of weight. Searched parameter values were $[0.0005, 0.001, 0.0025, 0.005, 0.01]$ for xenon.

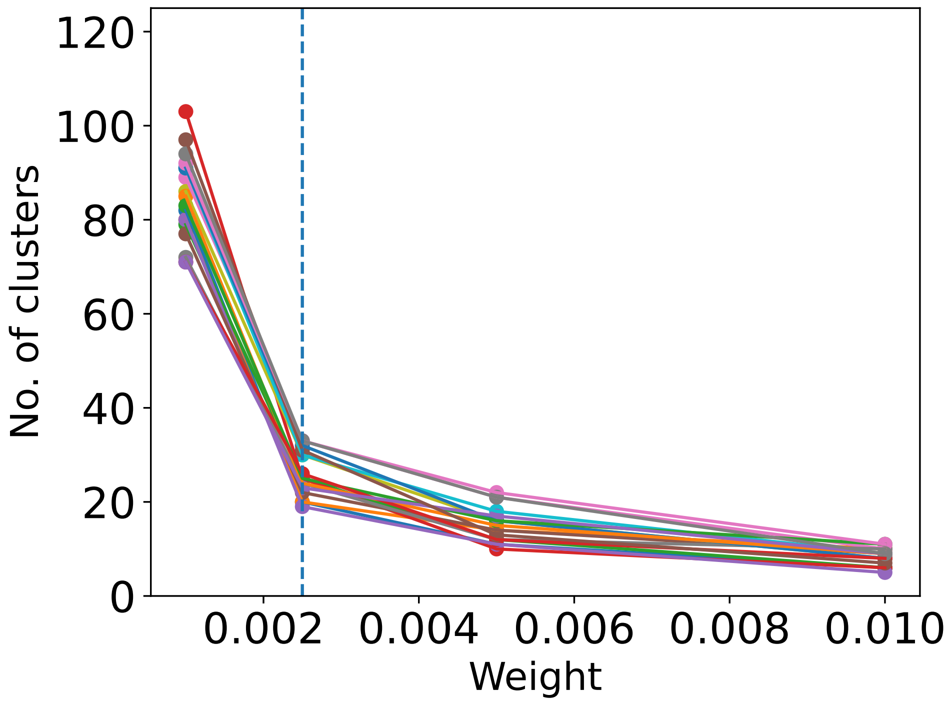


Figure S4: Parameter search for *min_clust_size*. Each coloured line indicates the dependency on weight for a simulation system. The dashed line points to the optimal value of weight. Searched parameter values were $[0.001, 0.0025, 0.005, 0.01]$ for benzene.

Table S3: Parameters of HDBSCAN. These are default values except for *min_clust_size*

| Parameter | Value |
|---|---|
| algorithm | 'best' |
| alpha | 1.0 |
| approx_min_span_tree | True |
| gen_min_span_tree | False |
| leaf_size | 40 |
| memory | None |
| metric | 'euclidean' |
| min_samples | None |
| p | None |

#### Reproducibility of xenon-binding events

To verify the reproducibility of the binding of xenon to known xenon-binding sites, the free energy iso-surface from each of the five independent MD simulations was visually compared to the known crystallographic xenon-binding sites (Figure S5). A simulation was considered successful if a stable, high-density xenon site overlapped with at least one of the known binding sites, as indicated by a check mark in Figure S5.

The results indicate that the binding events are highly reproducible for most systems. The success rate out of five simulations for each system was as follows: 4/5 simulations identified the known xenon-binding sites (Figure S5A), 1/5 simulations did (Figure S5B), 4/5 simulations did (Figure S5C), 4/5 simulations did (Figure S5D), 5/5 simulations did (Figure S5E), 5/5 simulations did (Figure S5F), and 5/5 simulations did (Figure S5G). This consistency across independent trajectories demonstrates that the observed xenon binding is a converged feature of the simulations.

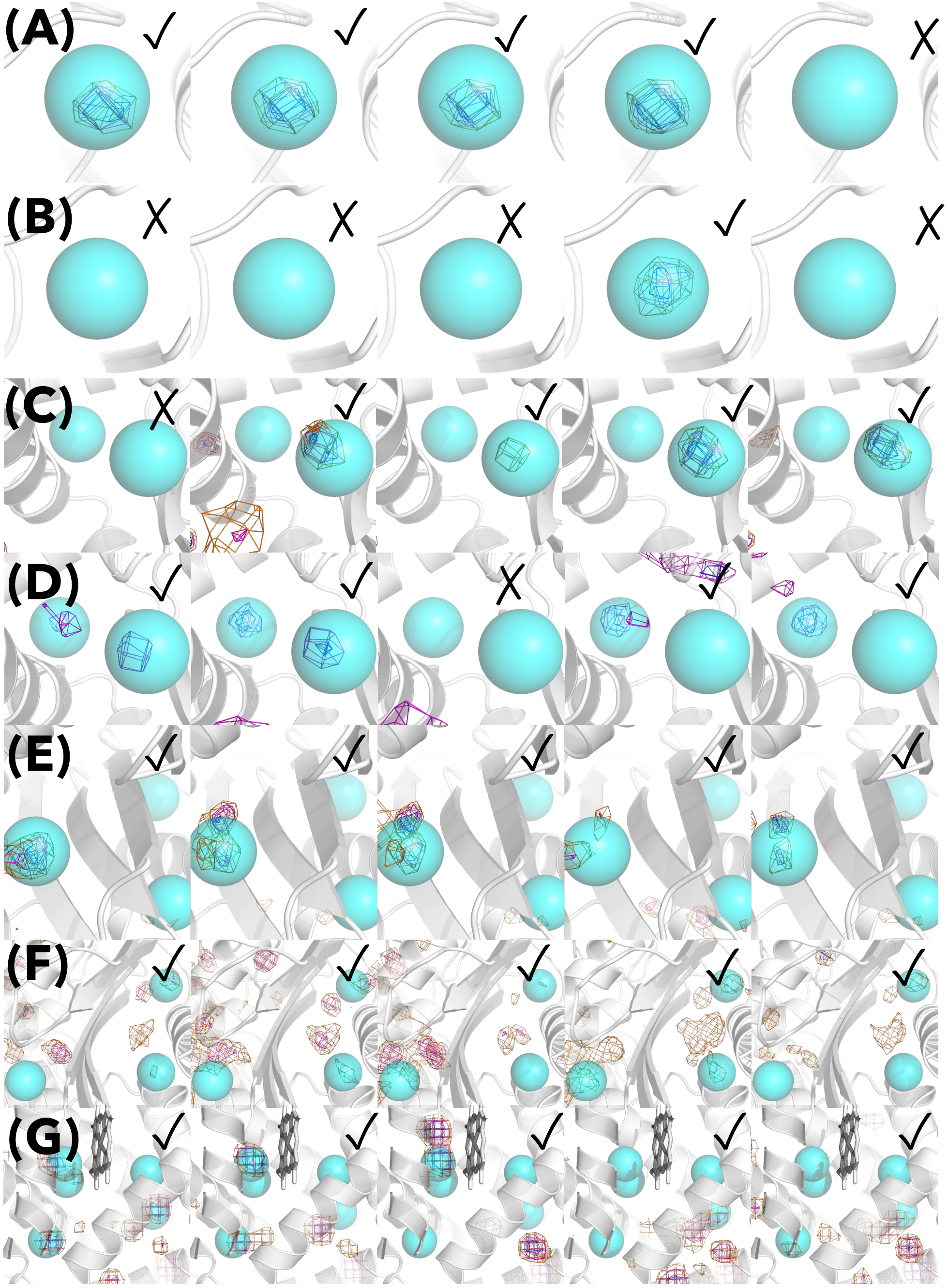


Figure S5: Free-energy iso-surface of xenon in each system. The corresponding PDB IDs are (A) 1c3l, (B) 1c1m, (C) 1c10, (D) 1c10(400K), (E) 5hw1, (F) 5hw1(400K), (G) 4nxa: The iso-surface $F(\boldsymbol{x})/k_\mathrm{B}T$ was coloured by the free energy value 3.0 for orange, 2.0 for magenta, and 1.0 for blue. If a high-density xenon site overlapped with at least one of the known xenon-binding sites, a check mark (✓) is added in each figure, otherwise a cross mark.

### Root mean squared deviation for xenon-protein complexes

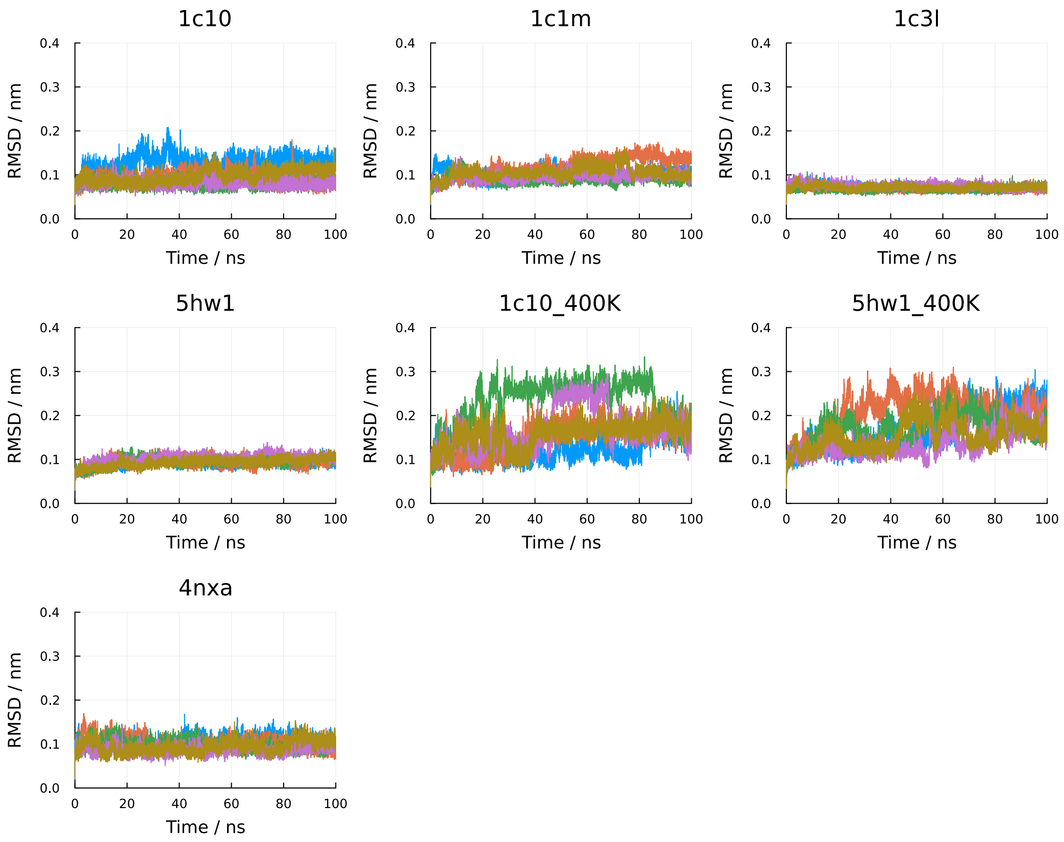


Figure S6: Five trajectories of RMSD for each target proteins with xenon atoms at 300 K and 400 K. The header in each plot indicates the PDB ID with or without the suffix of temperature value. No suffix means the results at 300 K

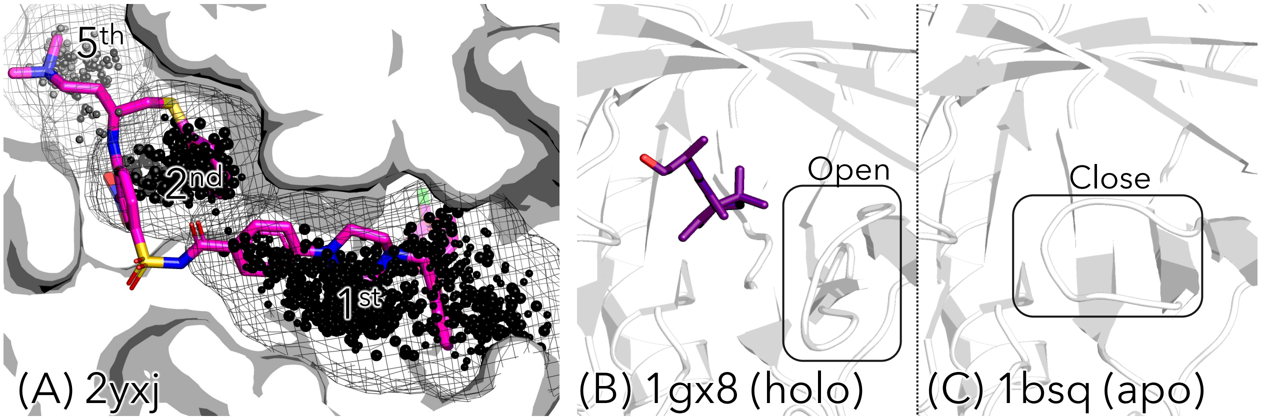


Figure S7: Success and failed cases of CB sites identification by benzene molecule. (A): A success case (PDB ID: 2yxj). The black spheres indicate the centre-of-mass point of a benzene molecule, which are mapped onto the holo structure 2yxj. The benzene molecules correctly identified the ligand binding sites including CB sites. (B) and (C): A failed case.

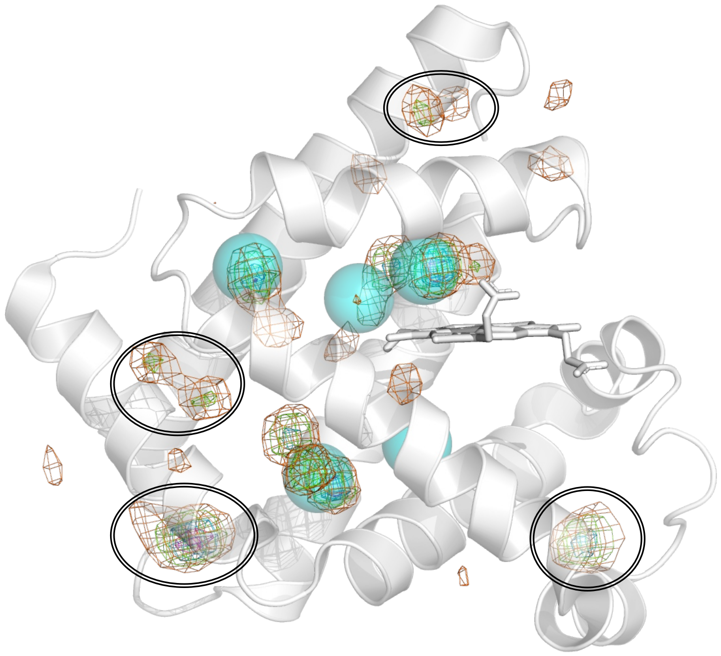

Figure S8: Overall picture of free-energy iso-surface of xenon in the myoglobin system. The iso-surface is mapped onto the myoglobin X-ray structure (PDB ID : 4nxa). The iso-surface $F(\boldsymbol{x})/k_{\mathrm{B}}T$ was coloured by the free energy value 3.0 for orange, 2.0 for magenta, and 1.0 for blue. The four double-line circles indicate xenon-binding sites that are not experimentally verified.

Table S4: Identification of **apparent binding sites** by **xenons** depending on the top $n$ clusters. The bold IDs indicate that apparent binding sites were not identified.

| PDBID(Apo) | Cluster threshold | | | | | | | | | |
|---|---|---|---|---|---|---|---|---|---|---|
| | 1 | $\leq 2$ | $\leq 3$ | $\leq 4$ | $\leq 5$ | $\leq 6$ | $\leq 7$ | $\leq 8$ | $\leq 9$ | $\leq 10$ |
| **3cj0** | False | False | False | False | False | False | False | False | False | False |
| 1uk3 | False | False | True | True | True | True | True | True | True | True |
| 1k5h | True | True | True | True | True | True | True | True | True | True |
| 1w50 | True | True | True | True | True | True | True | True | True | True |
| 1m47 | False | False | True | True | True | True | True | True | True | True |
| **1nep** | False | False | False | False | False | False | False | False | False | False |
| 1maz | True | True | True | True | True | True | True | True | True | True |
| 2wgb | False | False | False | False | False | False | False | False | True | True |
| 1jbu | False | False | False | False | False | False | False | False | True | True |
| 1bsq | True | True | True | True | True | True | True | True | True | True |
| 2ohg | False | False | True | True | True | True | True | True | True | True |
| 1b6b | False | True | True | True | True | True | True | True | True | True |
| **1hag** | False | False | False | False | False | False | False | False | False | False |
| 3f74 | False | False | True | True | True | True | True | True | True | True |
| 1fvr | False | False | False | True | True | True | True | True | True | True |
| 5j2v | False | True | True | True | True | True | True | True | True | True |

Table S5: Identification of **apparent binding sites** by **benzenes** depending on the top $n$ clusters. The bold IDs indicate that apparent binding sites were not identified.

| PDBID(Apo) | Cluster threshold | | | | | | | | | |
|---|---|---|---|---|---|---|---|---|---|---|
| | 1 | $\leq 2$ | $\leq 3$ | $\leq 4$ | $\leq 5$ | $\leq 6$ | $\leq 7$ | $\leq 8$ | $\leq 9$ | $\leq 10$ |
| **3cj0** | False | False | False | False | False | False | False | False | False | False |
| 1uk3 | False | False | False | False | False | False | False | True | True | True |
| **1k5h** | False | False | False | False | False | False | False | False | False | False |
| 1w50 | False | False | False | True | True | True | True | True | True | True |
| 1m47 | False | False | False | False | False | False | True | True | True | True |
| 1nep | False | True | True | True | True | True | True | True | True | True |
| 1maz | True | True | True | True | True | True | True | True | True | True |
| 2wgb | False | False | False | False | False | False | False | False | False | True |
| 1jbu | False | False | True | True | True | True | True | True | True | True |
| **1bsq** | False | False | False | False | False | False | False | False | False | False |
| **2ohg** | False | False | False | False | False | False | False | False | False | False |
| 1b6b | True | True | True | True | True | True | True | True | True | True |
| **1hag** | False | False | False | False | False | False | False | False | False | False |
| **3f74** | False | False | False | False | False | False | False | False | False | False |
| 1fvr | False | False | False | True | True | True | True | True | True | True |
| **5j2v** | False | False | False | False | False | False | False | False | False | False |

Table S6: Identification of **cryptic binding sites** by **xenons** depending on the top $n$ clusters. The bold IDs indicate that cryptic binding sites were not identified.

| PDBID(Apo) | Cluster threshold | | | | | | | | | |
|---|---|---|---|---|---|---|---|---|---|---|
| | 1 | $\leq 2$ | $\leq 3$ | $\leq 4$ | $\leq 5$ | $\leq 6$ | $\leq 7$ | $\leq 8$ | $\leq 9$ | $\leq 10$ |
| 1bsq | False | True | True | True | True | True | True | True | True | True |
| **2ohg** | False | False | False | False | False | False | False | False | False | False |
| 1b6b | False | False | True | True | True | True | True | True | True | True |
| **1hag** | False | False | False | False | False | False | False | False | False | False |
| 3f74 | True | True | True | True | True | True | True | True | True | True |
| 5j2v | False | True | True | True | True | True | True | True | True | True |
| 1fvr | False | False | False | True | True | True | True | True | True | True |
| 1uk3 | False | False | True | True | True | True | True | True | True | True |
| **3cj0** | False | False | False | False | False | False | False | False | False | False |
| 1k5h | True | True | True | True | True | True | True | True | True | True |
| **1w50** | False | False | False | False | False | False | False | False | False | False |
| 1m47 | False | False | True | True | True | True | True | True | True | True |
| 1nep | True | True | True | True | True | True | True | True | True | True |
| 1maz | True | True | True | True | True | True | True | True | True | True |
| 2wgb | False | False | False | False | False | False | False | False | True | True |
| 1jbu | False | True | True | True | True | True | True | True | True | True |

Table S7: Identification of **cryptic binding sites** by **benzenes** depending on the top $n$ clusters. The bold IDs indicate that cryptic binding sites were not identified.

| PDBID(Apo) | Cluster threshold | | | | | | | | | |
|---|---|---|---|---|---|---|---|---|---|---|
| | 1 | $\leq 2$ | $\leq 3$ | $\leq 4$ | $\leq 5$ | $\leq 6$ | $\leq 7$ | $\leq 8$ | $\leq 9$ | $\leq 10$ |
| **1bsq** | False | False | False | False | False | False | False | False | False | False |
| **2ohg** | False | False | False | False | False | False | False | False | False | False |
| 1b6b | True | True | True | True | True | True | True | True | True | True |
| **1hag** | False | False | False | False | False | False | False | False | False | False |
| 3f74 | True | True | True | True | True | True | True | True | True | True |
| **5j2v** | False | False | False | False | False | False | False | False | False | False |
| **1fvr** | False | False | False | False | False | False | False | False | False | False |
| **1uk3** | False | False | False | False | False | False | False | False | False | False |
| **3cj0** | False | False | False | False | False | False | False | False | False | False |
| **1k5h** | False | False | False | False | False | False | False | False | False | False |
| **1w50** | False | False | False | False | False | False | False | False | False | False |
| 1m47 | False | False | False | False | False | False | False | False | True | True |
| 1nep | True | True | True | True | True | True | True | True | True | True |
| 1maz | True | True | True | True | True | True | True | True | True | True |
| **2wgb** | False | False | False | False | False | False | False | False | False | False |
| 1jbu | True | True | True | True | True | True | True | True | True | True |
